# Visual association cortex links cues with conjunctions of reward and locomotor contexts

**DOI:** 10.1101/2021.08.07.453879

**Authors:** Kelly L. McGuire, Oren Amsalem, Arthur U. Sugden, Rohan N. Ramesh, Christian R. Burgess, Mark L. Andermann

## Abstract

Postrhinal cortex (POR) and neighboring lateral visual association areas are necessary for identifying objects and interpreting them in specific contexts, but how POR neurons encode the same object across contexts remains unclear. Here, we imaged excitatory neurons in mouse POR across tens of days throughout initial cue-reward learning and reversal learning. As such, neurons were tracked across sessions/trials where the same cue was rewarded or unrewarded, during both locomotor and stationary contexts. Surprisingly, a large class of POR neurons were minimally cue-driven prior to learning. After learning, distinct clusters within this class responded selectively to a given cue when presented in a specific conjunction of reward and locomotion contexts. In addition, another class involved clusters of neurons whose cue responses were more transient, insensitive to reward learning, and adapted over thousands of presentations. These two classes of POR neurons may support context-dependent interpretation and context-independent identification of sensory cues.

## INTRODUCTION

Our reactions to objects change with experience. Yet even for familiar objects, the correct actions to take and outcomes to expect can differ across diverse spatial, reward, and internal contexts in a manner that evolves over time.

Physiology and silencing studies suggest that rodent postrhinal cortex (POR; Beaudin et al., 2013; Wang and Burkhalter, 2007) and adjacent areas are important for associating visual cues with rewards (Burgess et al., 2016; Ramesh et al., 2018; Sugden et al., 2020) or punishments (Sacco and Sacchetti, 2010), and for associating visual objects with the spatial contexts in which they occurred (Burwell et al., 2004; Furtak et al., 2012; Norman and Eacott, 2005). In addition, POR exhibits characteristics of early visual cortical areas, such as retinotopic organization and brisk visual responses in naïve animals (Burgess et al., 2016; Beltramo and Scanziani, 2019).

Identifying the appropriate meaning of visual cues in specific reward and spatial contexts likely involves inputs to POR from neurons encoding both cues and rewards in basolateral amygdala (Burgess et al., 2016) and from neurons encoding spatial and non-spatial contexts in entorhinal cortex and prefrontal cortex (Burwell and Amaral, 1998; Reinert et al., 2021; Wilson et al., 2013). Studies of primary visual cortex and other cortices in naïve animals have demonstrated strong (and mostly positive) influences of “internal” contexts such as whether the animal is running or stationary when the cue appears (Andermann et al., 2011; Erisken et al., 2014; Niell and Stryker, 2010; Pakan et al., 2016; Saleem et al., 2013). Recent work has also shown that the transition between stationary and locomotor contexts causes a rapid and bidirectional reconfiguration of the dialogue between individual V1 neurons and distinct cortical regions (Clancy et al., 2019), suggesting a more nuanced role for various contextual influences from other brain areas across distinct locomotor states.

While early electrophysiology experiments showed differences in perirhinal and postrhinal cortex responses to objects in different spatial contexts (Furtak et al., 2012; Rolls et al., 2005), little is known about how the network of neurons in POR and nearby areas encode the same cues throughout many weeks of experience and across a variety of contexts. Does the same common set of neurons exhibit responses to a cue throughout learning but with changing magnitude and dynamics? Or alternatively, are previously silent neurons recruited to encode novel memories of cues in specific contexts (Cai et al., 2016; Ramesh et al., 2018; Rashid et al., 2016)?

To address these questions, we used longitudinal two-photon calcium imaging in transgenic mice to track visual cue responses in thousands of POR neurons across tens of training sessions from the naive state through initial association learning and subsequent re-learning following a reversal in cue-outcome contingencies. We took an unbiased approach to visualizing and clustering populations of neurons with similar changes in cue responses across stages of learning using tensor component analysis (TCA; Williams et al., 2018). Using this paradigm, we observed multiple groups of neurons with shared within-stage and across-stage response dynamics, revealing a division of labor between clusters of neurons that encoded a given cue (i) only when the cue was rewarded, (ii) only when the cue was not rewarded, or (iii) in a manner that was independent of reward context. Each of these clusters contained roughly balanced subgroups of neurons that were strongly positively or negatively modulated by locomotion across stages of learning. We also found other groups of neurons that responded to cue *offsets* in either a reward context-dependent or context-independent manner. Taken together, these findings suggest that POR contains intermingled groups of neurons that encode sensory cues either independent of context (potentially useful for invariant object recognition) or in specific conjunctions of contexts (i.e., only during locomotion or when stationary, and only when the cue is rewarded or unrewarded). We suggest that learned encoding of cues in specific conjunctions of contexts facilitates readout of cue-context associations that form the basis for context-dependent selective attention and action selection.

## RESULTS

### Longitudinal imaging of hundreds of neurons throughout associative learning and reversal relearning

We investigated cue-outcome learning in food-restricted, head-restrained mice using the same Go/NoGo visual discrimination task that we have described previously (Fig. 1A, B; Burgess et al., 2016; Livneh et al., 2017; Ramesh et al., 2018). We examined changes in visual cue responses in excitatory neurons in layer 2/3 of postrhinal cortex and adjacent regions of lateral visual association cortex (hereafter referred to as POR) using chronic two-photon calcium imaging in transgenic mice expressing GCaMP6f (Fig. 1C-E; Sugden et al., 2020). The imaging field of view was centered on area POR using retinotopic mapping (Burgess et al., 2016). The use of transgenic mice enabled more stable tracking of 7,824 neurons across ∼10,000 trials (median number of trials per mouse: 10,838 across 7 mice, range: 5,166-12,813 trials; median number of imaging sessions per mouse: 25, range: 11-33; Fig. 1F, bottom), starting from the naïve state (in 4/7 mice) or starting from initial stages of cue-outcome association (3/7 mice). In particular, we focused our analysis on cue responses across five stages of increasing mastery of the task (“initial learning”), as well as during another five stages of increasingly improved performance following a switch in cue-outcome contingencies (“reversal learning”, Figure 1F, Fig. S1A; Ramesh et al., 2018). While learning did progress in a largely monotonic fashion (Fig. S1B), these stages were binned by equally spaced increments in task performance (discriminability, d’; Poort et al., 2015) rather than in time.

**Figure 1.**
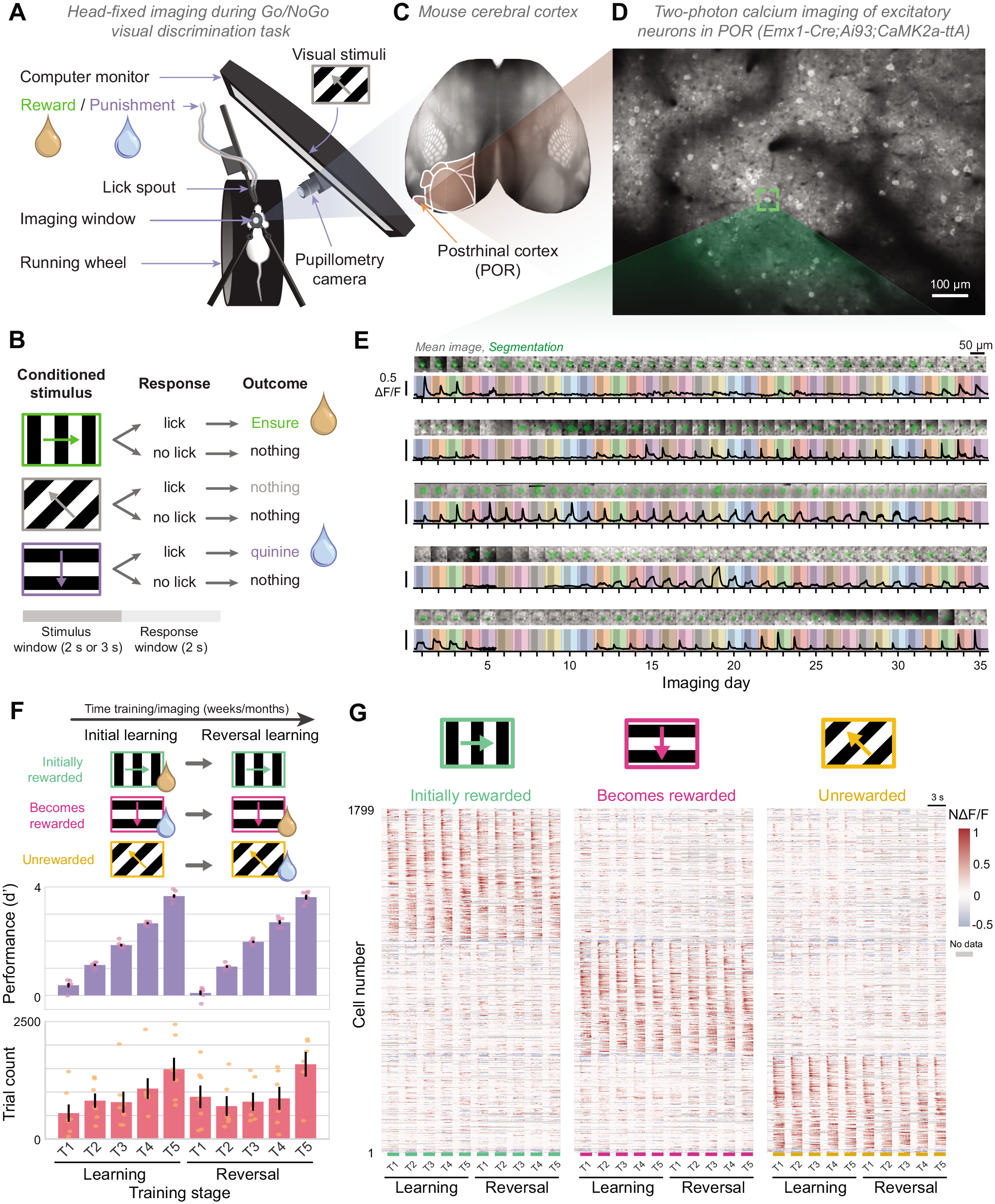
Longitudinal imaging of hundreds of neurons throughout visual association learning and relearning. A. Head-restrained setup for visual stimulation, delivery of reward or punishment, and two-photon calcium imaging. B. Animals were trained on a three-orientation operant Go/NoGo visual discrimination task. C. Mouse cerebral cortex, highlighting visual areas (pink) and postrhinal visual association cortex (POR). Image credit: Allen Institute. D. Example field of view centered on POR (depth: 149 μm). For each mouse, cells from one field of view were aligned and tracked over weeks and months of imaging during learning. E. Mean cell image (grayscale) and segmentation (green) for five example cells, together with average stimulus-evoked response per day across 35 days of imaging. ΔF/F: fractional change in fluorescence. Colored shading indicates different days. Darker regions denote stimulus period (3 s). F. Initial learning and acquisition of the task was followed by a change in cue-outcome contingencies during reversal learning (top). Stages of learning across training were divided into evenly spaced bins (middle) according to behavioral performance (d’; Stages T1 learning, …, T5 learning, T1 reversal, …, T5 reversal). This approach allowed averaging of responses across many hundreds of cue presentations per stage (bottom). In middle and bottom plots, dots indicate individual mice, bars indicate averages across mice. G. Heatmap of mean responses across trials in each of 10 stages of learning for all cue-driven cells (n = 1799 cells from 7 mice). Unless otherwise specified, responses of each cell are normalized to the maximum response across all stages and cues. Thus, a normalized unit of responsiveness (NΔF/F) is shown. Cells are sorted into those that respond preferentially to each cue, and further sorted by peak response latency (see Methods). See also Figure S1.

We found that, in general, the temporal dynamics (e.g., sustainedness) and cue preference of each neuron’s sensory responses were stable in successive stages of learning and relearning. This is clear in Fig. 1G, which shows the response time course of each significantly cue-driven neuron (1799 cue-driven neurons out of 7824 neurons that were tracked across multiple days from 7 mice, see Methods), averaged across all trials in each stage of learning for cells which had a significant response in at least one stage. However, individual neurons showed a range of response dynamics. When we sorted neurons responsive to each cue by their peak response latency, we found that neurons with short peak latencies tended to have a transient response, while neurons with longer peak latencies tended to have a more sustained response throughout the stimulus period (Fig. 1G; see also Fig S1C and Fig. 3, below).

### Evolution of responses in distinct clusters of neurons across learning and relearning

Having established an experimental setup for longitudinal tracking of visual cue responses beginning in naïve mice or early learning and extending through task mastery and subsequent relearning, we asked whether these data might reveal functional classes of neurons that could not be ascertained in previous studies. Two example neurons in Fig. 2A illustrate the diverse changes we observed in both response magnitude and response shape across stages of learning. Accordingly, we sought a dimensionality reduction method that could classify neurons with distinct response dynamics (possibly reflecting varying influences of different sources of input) into different classes.

**Figure 2.**
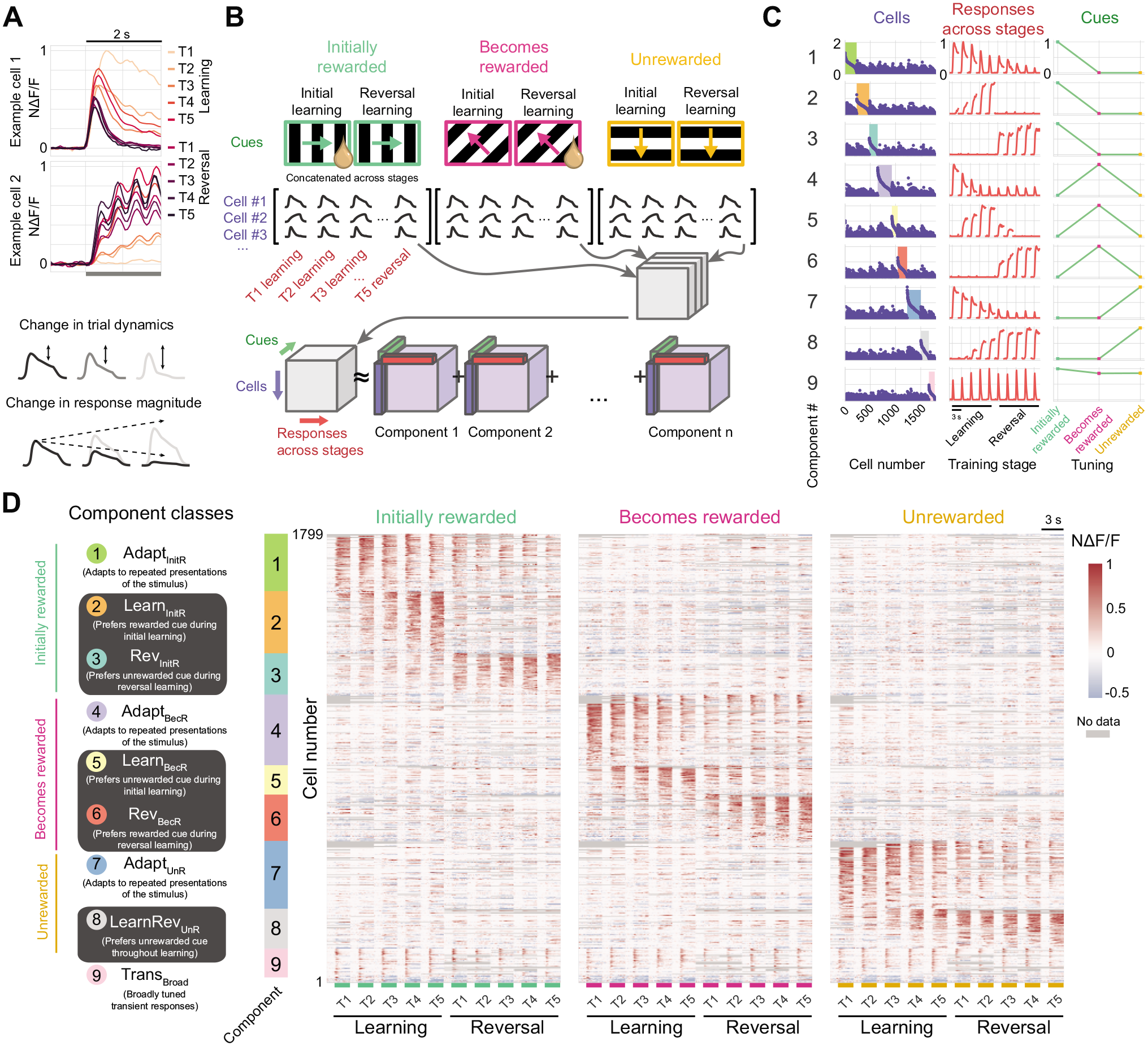
Evolution of responses in distinct classes of neurons across learning and relearning. A. Top: example traces from two neurons showing changes in responses across learning (normalized to peak response across stages). The activity of one neuron also tracked the 2 Hz stimulus frequency. Bottom: schematic demonstrating two possible timescales of neuronal response changes across learning. Individual cells could show within-stage changes in response shape and/or across-stage changes in response magnitude. B. Schematic of data organization for tensor component analysis (TCA). Average response time courses (1-s baseline, 2-s stimulus response) for each training stage were concatenated, resulting in a single time course of responses to each of the 3 cues, for each cue-driven cell (n = 1799 cells). TCA finds components of activity represented by low dimensional factors corresponding to groups of cells (cell factors), patterns of activity across stages of learning (temporal factors), and relative magnitude of responses to each visual cue (tuning factors). C. Cell factors, temporal factors, and tuning factors are shown for all nine components of a rank 9 TCA model. Different tensor components capture distinct population dynamics both within trials and across stages of learning. Note the clear change in response magnitude in some components following reversal. Colored bars (left) highlight cells whose weight was highest for that component versus all others. D. Mean responses during each training stage (Fig. 1G), sorted as in C, according to each cell’s affiliation with a component from this rank 9 model. Component numbers from C are indicated in colored circles. This classification highlights clusters of cells whose responses either adapt across training, as well as other clusters of cells (highlight with dark gray background in left column) that did not adapt across learning but instead showed responses specific to when a cue was either rewarded or unrewarded. The latter clusters exhibit responses that peak following successful initial learning or following successful reversal learning. A final cluster showed transient and broadly tuned responses that remain constant across stages. A small set of cells (bottom rows; no associated colorbar) could not be well classified (see Methods). Cells were first sorted by cue preference (see tuning factors in C) and then by cell factor weight. See also Figure S2.

To account for changes in both the overall magnitude of the cue response and the shape of the cue response across stages of learning, as well as for differences in cue preferences across neurons, we employed a modified implementation of tensor component analysis (TCA; Williams et al., 2018) (Fig. 2B, see Methods). We first created a three-dimensional matrix of response dynamics for all cue-driven neurons (i.e., a tensor with dimensions: [neurons] x [peak-normalized concatenation of stage-averaged time courses for each neuron] x [“Initially rewarded” cue, “Becomes rewarded” cue, “Unrewarded” cue]). Tensor component analysis allows optimal decomposition of this 3D dataset into multiple 3D components. Each component is the outer product of (i) a vector of relative weights, or contributions, for each cell, (ii) a time course reflecting both within-trial and across-stage temporal dynamics (a “temporal factor”), and (iii) a tuning curve of response magnitudes across the three cues (Fig. 2B). Given the sharp tuning of most neurons for one of the three cues, and the diversity in the response shape and across-stage changes in response magnitude of various neurons (Figs. 1G, 2A), we fit a range of TCA models that differed in the number of components (i.e., models of different ranks) in order to empirically determine a reasonable number of clusters of neurons. We also ran multiple iterations for each model rank to ensure model stability (Fig. S2). We found that these models could efficiently capture key features of this large dataset across ranks (Fig. S2A-C) and yielded qualitatively similar results across iterations at a given rank (Fig. S2C, D). We observed four common classes of components in this example model and across a range of models varying in the number of components (e.g., from 6-12, Fig. S2C top, E). This is illustrated by the nine components from an example rank nine model shown in Fig. 2C (rows are sorted by preferred cue). First, we observed three components — one tuned to each cue — whose temporal factors exhibited partial response adaptation during the cue period, and a slow, progressive attenuation of net responses across stages that did not abruptly change across the reversal (Fig. 2C, 1^st^, 4^th^ and 7^th^ rows; Fig. S2A, C, E). We also observed components that showed sustained responses to one of the three cues (indicated by dark gray shading in Fig. 2D, left), either during later stages of initial learning (Fig. 2C, 2^nd^ and 5^th^ rows; Fig. S2A, C, E), during later stages of reversal learning (Fig. 2C, 3^rd^ and 6^th^ rows; Fig. S2A, C, E), or throughout later stages of initial learning and all stages of reversal learning (Fig. 2C, 8^th^ row; Fig. S2A, C, E). Specifically, we observed five sustained components: one responsive to the rewarded cue before reversal, one responsive to the newly rewarded cue after reversal, and one responsive to each of the three cues when that cue was unrewarded. We also often observed a ninth component with short-latency, transient responses that did not change across stages of learning and were not sharply tuned to a given cue (Fig. 2C, bottom row; Fig. S2C, E). Given that a range of models and model ranks (Fig. S2) yielded components with qualitatively similar dynamics to this nine-component model in Fig. 2C, we decided to further characterize groups of neurons using this nine-component model.

We classified neurons by assigning each cell to the component that best described that cell’s activity, defined as the component with the highest rescaled cell factor weight (see Methods and shaded areas in Fig. 2C, left). In this way, we could re-sort the plot of each cell’s response time courses (Fig. 1G) according to the cell’s assigned cluster (Fig. 2D; using the same sorting of cells as in Fig. 2C, left column). This revealed that the within-trial and across-stage dynamics of each of the nine components was faithfully reflected in the dynamics of the associated cluster of constituent cells, each tracked over many stages of learning and reversal learning. By tracking the entire learning process, we could classify neurons based on features that derive in large part from common changes across many sessions during learning and reversal learning, but which would not be evident if only evaluating single sessions or stages of learning (see individual columns in Fig. 1G). Averaging the responses across cells in each cluster from each mouse confirmed that the distinct across-stage response dynamics in each of the nine functional clusters was not due to pooling of functionally distinct sets of cells across different mice (Fig. 3A).

**Figure 3.**
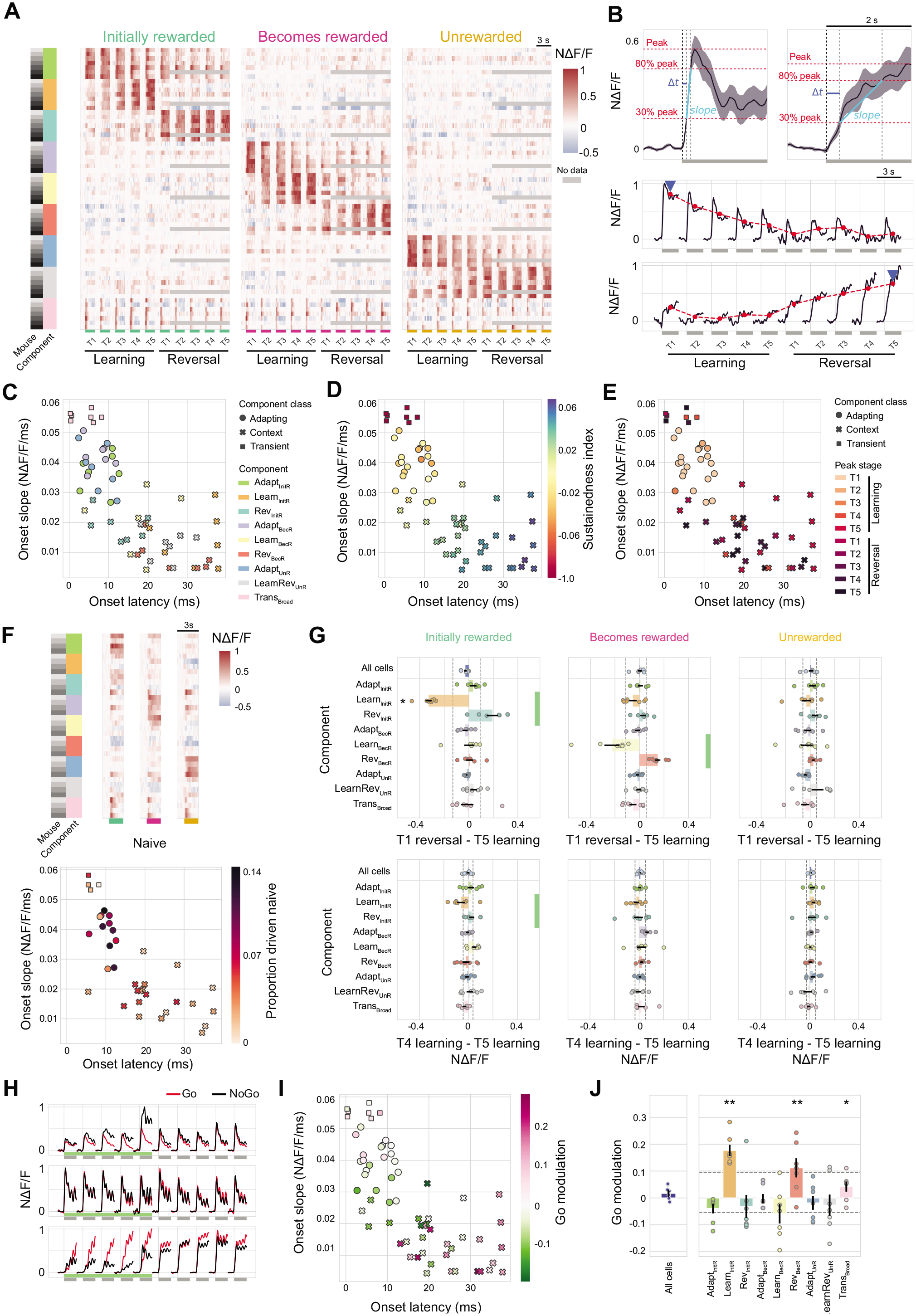
Cells with distinct within-trial and across-learning dynamics show differential sensitivity to reward context. A. Average response traces across cells clustered by component for each mouse. Each row represents the average response across cells associated with a component for a single mouse. Cell averages for each mouse produced qualitatively similar responses. Components are sorted as in Fig. 2D. B. Top: estimation of onset latency and onset slope for two example response traces (averaged across stages). Onset latency (blue line) was calculated as the first crossing of a threshold (30% of peak response) following stimulus onset. Onset slope (cyan line) was calculated between 30% and 80% threshold crossings (red lines). Bottom: estimation of mean response per stage (red line) and of maximally responsive stage (i.e., “Peak” stage; blue arrowhead). C. Scatter of onset latency versus onset slope. Each data point represents one mouse and component. The shape of each data point denotes whether the component was considered an “Adapting,” “Context,” or “Transient” component. Color denotes component cluster. D. Same as C, except that color indicates the “sustainedness” index of the average stimulus response of that component cluster. E. Same as C, except that color indicates the peak stage of the response across learning. F. Top: in a subset of mice (n = 4), we imaged before training began (i.e., naïve passive viewing) to further characterize cell dynamics across learning. Cell masks from naive sessions were aligned to those during learning, and mean responses across all naïve trials were calculated as in A. Bottom: the proportion of cells driven during imaging in naïve mice was plotted for each component using the same axes as in C. One data point per mouse and component. G. Difference in normalized response magnitude for average traces of each component (cf. red dots in B, bottom) between two neighboring stages of training that either straddled the reversal (top; T5 learning versus T1 reversal) or that preceded the reversal (bottom; T4 learning versus T5 learning). Green bars indicate component clusters whose preferred cue is rewarded during at least one of the two stages. *p < 0.05, **p < 0.001; bootstrap test over stages, Bonferroni corrected across components (including “All cells”). Dashed line: 95% confidence interval estimated using bootstrapping across components. Error bars: SEM across mice. H. Average response to the initially rewarded cue for three example cells, plotted separately for Go trials (red) or NoGo trials (black) for each stage of learning. Green bar indicates stages for which the visual cue is associated with reward. Gray bar indicates the 2 s after visual stimulus onset. I. Same as C, except that color indicates the average Go modulation of cells in each component (Go modulation = [Mean Go resp. - Mean NoGo resp.]). J. Difference in neural response for Go trials versus NoGo trials, averaged across cells in each component cluster and mouse. This difference was also calculated after averaging trials across all cells from all component clusters in each mouse (left, blue). *p < 0.05, **p < 0.001; bootstrap test over Go/NoGo trial averages, Bonferroni corrected for components (including “All cells”). Dashed line: 95% confidence interval estimated using bootstrapping across components. Error bars: SEM across mice. See also Figure S3.

While across-stage dynamics and cue selectivity were clearly central to the fitting and classification of these functional clusters, we also found that these clusters of neurons differed in their within-trial dynamics. Specifically, we averaged response time courses across stages and across cells in each cluster, and characterized the onset latency and slope of the rising phase of the response (onset slope) (Fig. 3B-C). We also characterized the response “sustainedness” of each cluster (Fig. 3D, S3A), and the stage of learning where the response in each cluster was maximal. A scatter plot of onset latency versus onset slope for each of the nine component clusters per mouse delineated three broad classes of neurons (Fig. 2D, 3C, S3B). The first class was composed of a single cluster of neurons that exhibited broadly tuned, fast-transient responses (“Trans_Broad_” component, Fig. 2D, 3A) with the shortest onset latencies and steepest onset slopes, possibly reflecting strong bottom-up input (Fig. 3C, S3B). This cluster also had the least sustained responses across the cue period (Fig. 3D) and showed inconsistent and weak preferences for any given stage of learning (Fig. 3E, S3C), reflecting similar response magnitudes across stages (Fig. 3A). A second class was composed of three clusters of adapting cells that selectively responded to the cue that was initially rewarded (“Adapt_InitR_” component), the cue that became rewarded (“Adapt_BecR_” component), or the cue that remained unrewarded (“Adapt_UnR_” component, associated with no outcome or with avoidable quinine delivery, a mild punishment). Adapting clusters all showed intermediate onset latencies and onset slopes across mice (Fig. 3C, S3B), and intermediate response sustainedness, consistent with partial response adaptation across the cue period (Fig. 3D). Adapting clusters also showed preferred responses to the initial stages of learning, with progressively weaker responses across successive stages (Fig. 3A, 3E, S3C). The remaining class was composed of five clusters of neurons that all showed longer onset latencies and less steep onset slopes (Fig. 3C, S3B) and more gradual buildup of activity during the cue period (Fig. 3B). These clusters also showed preferences to later stages in learning (Fig. 3E, S3C), either stages of high performance during initial learning (“Learn_InitR_”, “Learn_BecR_” components), during reversal learning (“Rev_InitR_”, “Rev_BecR_” components), or across both initial and reversal learning (“LearnRev_UnR_” component). For these five clusters, the emergence of peak responses at later stages of learning were paralleled by increasing numbers of significantly driven cells within each cluster at these later stages (Fig. S3D). Thus, in contrast to fast-transient cells which responded throughout learning (consistent with Ramesh et al., 2018), these “reward-context-dependent” clusters predominantly responded to only one cue — but only in a single context in which that cue was unrewarded or rewarded — and otherwise were mostly silent.

The preferences of adapting clusters of cells for early stages of learning, and their continued responsivity across learning, suggested that they encode visual stimuli regardless of the associated actions/outcome contexts. Additional evidence supporting this idea came from additional recordings in a subset of these mice *prior* to any pairing of outcomes with these cues (i.e., in “naïve” mice). Indeed, prior to any training, adapting clusters of cells showed more consistent responses to their preferred cues than the other cell clusters (n = 4 mice, Fig. 3F), accounting for over half of all driven cells (Fig. S3E). In contrast, clusters with cue responses that varied with reward context became more common in later stages of learning and relearning (Fig. S3D).

To more directly examine whether changes in the responses of these “reward-context-dependent” clusters over stages was indeed due to changes in cue meaning or instead due to a natural drift in neurons’ cue responsivity, we directly compared changes in the responses of each cluster of neurons across successive stages that did or didn’t involve a change in the association between cues and outcomes (see also Figs. 2D, 3A). When we considered changes in the average cue responses in each cluster between the last stage of initial learning and the following stage (which involved a switch in cue contingencies), we observed large, opposing changes in response magnitude in the clusters of neurons responsive to the cue that was initially rewarded, and in those responsive to the cue that became rewarded after reversal (Fig. 3G). These changes were consistent across mice. In contrast, no abrupt changes in response magnitude were observed for adapting clusters of cells sensitive to these two cues, consistent with a suggested role in context-*independent* sensory processing. Further, no abrupt changes in response magnitude were observed for any of the clusters of cells responsive to the third cue, which remained unrewarded both pre- and post-reversal and thus did not change its meaning. Similarly, no abrupt changes in response magnitude were observed for any clusters of neurons between the last two stages of initial learning, for which there was no change in cue meaning (Fig. 3G).

We reasoned that, if a cluster of neurons responded to a visual cue specifically in a learning stage where that cue predicted a reward, then this same cluster should respond more vigorously on those cue presentations that were followed by operant motor responses (Go trials) than for those not evoking a motor response (NoGo trials). Conversely, if a cluster of neurons responded to a cue specifically in learning stages where the cue did not predict a reward, then this same cluster should respond more vigorously on those cue presentations in which the animal does not anticipate reward delivery (NoGo trials). Example neurons with consistent biases to Go trials or to NoGo trials are shown in Fig. 3H. Across mice, clusters of neurons exhibiting a robust response bias to Go versus NoGo trials were typically those that selectively responded to a visual cue in stages when it was associated with a possible reward (Fig. 3I-J), while adapting clusters rarely showed a Go bias. NoGo response biases were observed in clusters of cells that only fired reliably during stages where the cue was not rewarded, as well as in some adapting clusters (Fig. 3I-J). Note that these Go or NoGo biases do not reflect pure motor-related responses. For example, clusters with Go biases only responded vigorously when one specific cue was rewarded, and did not show substantial responses in other stages when another cue evoked similar operant motor responses and reward delivery (Fig. 3A).

Taken together, the above findings suggest a division of labor of POR layer 2/3 excitatory neurons that respond to a given cue. Adapting clusters of neurons respond to the same cue beginning prior to or early on in learning, irrespective of the reward association, but gradually decrease their responses upon repeated presentation of the cue across dozens of sessions. Meanwhile, distinct clusters of neurons exist that respond to each sensory cue only in stages when it is associated with reward or only when it is associated with a lack of reward. These findings extend to reward contexts the notion that lateral visual association cortical regions play an important role in encoding objects in particular spatial contexts (Furtak et al., 2012; see Discussion).

### Distinct neurons encode cues in particular locomotor and task contexts

An additional context that shapes sensory processing is the locomotor state of the animal (see Discussion). In mice, excitatory neurons in mouse primary and secondary visual cortices exhibit visual responses that are strongly modulated by locomotion and associated arousal (mostly enhanced in primary visual cortex) (Andermann et al., 2011; Erisken et al., 2014; Niell and Stryker, 2010; Pakan et al., 2016; Saleem et al., 2013). Recent studies of ongoing activity in visual and retrosplenial cortical neurons have also shown that neurons dramatically change their functional correlations with local neurons and with distant brain areas during quiescence versus locomotion (Clancy et al., 2019). Therefore, we considered how locomotor state affects cue responses in POR. The aforementioned studies suggest that adapting neurons in POR, given their faithful, short-latency coding of visual stimuli, might also show consistent enhancement of visual responses during locomotion. Similarly, the relationship of locomotion with arousal (Vinck et al., 2015) might suggest that context-selective neurons encoding cues associated with motivationally salient rewards (but not neurons encoding neutral cues) would show further arousal-related enhancement of cue responses with locomotion. Instead, as shown below, we observed balanced sets of neurons that responded to a cue either when the mouse was running or when the mouse was stationary.

Example neurons in Figure 4A illustrate cases with either little effect of running or strong enhancement or suppression of visual responses with running. We separately considered responses of all cue-driven neurons, averaged separately across stationary trials or running trials in each stage of learning (Fig. 4B; cells preferring a given cue were sorted by their response sensitivity to locomotion). The mean responses of each cell during stationary versus running trials revealed distinct sets of neurons responsive to a specific cue only in a particular locomotor context, as well as neurons with less sensitivity to locomotion (Fig. 4C, left). Response biases to running or stationary states were fairly balanced across neurons (Fig. 4C, right). Strikingly, balanced sets of neurons that responded to a specific cue either during running or stationary states were observed for each of the nine clusters of neurons described in Figures 2-3 (Fig. 4D). Further, the mean response across neurons in each adapting cluster was similar in running or stationary states, and only showed a weak response bias to running states for other components (Fig. S4A). Individual neurons’ cue response biases to running states or to stationary states were stable and consistent across stages of learning (Fig. 4A, E-F, and S4B), suggesting that the influence of locomotor context is stable across learning.

**Figure 4.**
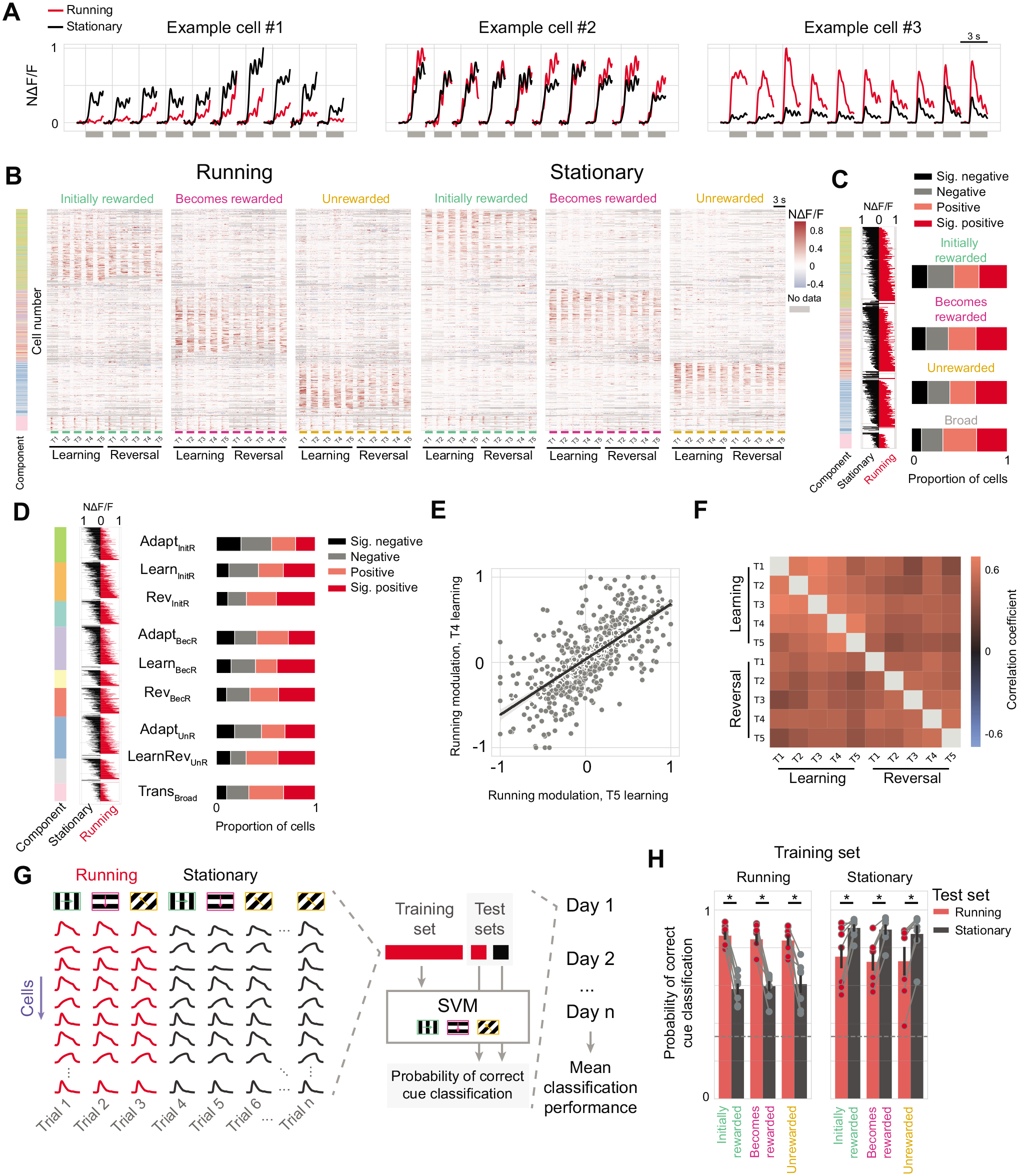
Distinct neurons encode cues in particular locomotor and task contexts. A. Three example neurons showing negative (left panel), neutral (middle panel), and positive (right panel) modulation by running across stages of learning. Modulation was consistent across all stages of initial learning and reversal learning. B-C. Cell response time courses per stage (B), averaged across running trials (left) and stationary trials (right). Colorbar indicates the component cluster to which each cell belongs, as defined in Fig. 2D. Cells are sorted by cue preference and then by running modulation index, estimated as the difference in normalized mean responses (pooled across stages) during running trials (red lines in panel C, left) versus stationary trials (black lines in C, left). We observed a substantial proportion of cells preferring each cue that were either significantly positively or negatively modulated by locomotion (C, right), as assessed using bootstrapping (see Methods). D. Same as C, but cells are first sorted by component cluster and then by running modulation. Note that each component contains sets of cells with significant positive or negative modulation by locomotion. E. Modulation by locomotion was similar for each cell when calculated during T5 or T4 stages of initial learning. Data were fit using linear regression. Pearson’s r: 0.68, p < 10^-79^. F. Pearson correlation of running modulation calculated between each pair of learning stages, illustrating consistency across weeks of recordings, including across reversal. G. Schematic of support vector machine (SVM) decoding analysis. An SVM was trained to classify which of the three visual stimuli were presented on a given trial from neural activity (e.g., using the mean population response vector from a given running and stationary trial). The SVM was trained using only locomotion trials or stationary trials, but tested on both locomotor contexts using held-out data. Models were trained and tested within each recording session, and the average performance (e.g., fraction of correctly classified trials) was averaged across sessions per mouse. H. Average classification performance per cue for the SVM trained on locomotion trials (left) or on stationary trials (right). One data point per mouse. Gray dashed line indicates chance decoding. *p < 0.05; one-way Wilcoxon signed-rank, Bonferroni corrected for cues. Error bars: SEM across mice. See also Figure S4.

Together, these results suggest that distinct sets of neurons exist in POR that selectively respond to specific cues in different *conjunctions* of task and locomotor contexts. As such, we hypothesized that knowledge of locomotor context should be important for downstream neurons to accurately decode population activity regarding cues. We confirmed this was the case using a support vector machine (SVM) classifier. Across mice, a classifier trained using running trials was far more accurate when tested on held-out running trials than on stationary trials, and vice versa (Fig. 4G-H).

### Average network activity is balanced across reward and locomotor contexts

Many previous studies of cortical activity during learning involved bulk recordings in humans and animal models (Harel et al., 2014; Hebart et al., 2018; Orsolic et al., 2021). Our chronic cellular imaging experiments reveal that different sets of context-specific neurons in POR respond to a cue at different stages of learning (Fig. 3D, 4A). Thus, we considered whether the *average* neural activity across this cortical region would also exhibit changes across learning. For each stage of learning, we averaged the cue-evoked response time courses of all neurons from Fig. 3D (without peak-normalizing each neuron’s activity across stages). The resulting population average time courses for the initially rewarded stimulus were remarkably stable across stages of learning, both in average magnitude and in within-trial response shape (Fig. 5A, *top*). In comparison, the average response across clusters of cells assigned to each component (as in Fig. 4A) showed dramatic differences in shape and in across-stage peak magnitudes (Fig. 5A, *bottom*). As shown in Fig. 5B, these differences in each component’s shape and in its responsivity across stages appeared to balance out, with the Adapt_InitR_ cells decreasing during stages where the Learn_InitR_ cells increases in magnitude. When this latter cluster of cells became less responsive following reversal, their responses were replaced by increased responses in Rev_InitR_ cells. A similarly balanced tradeoff in net responsivity by different context-dependent clusters of neurons was observed across individual mice (Fig. S5A-C).

**Figure 5.**
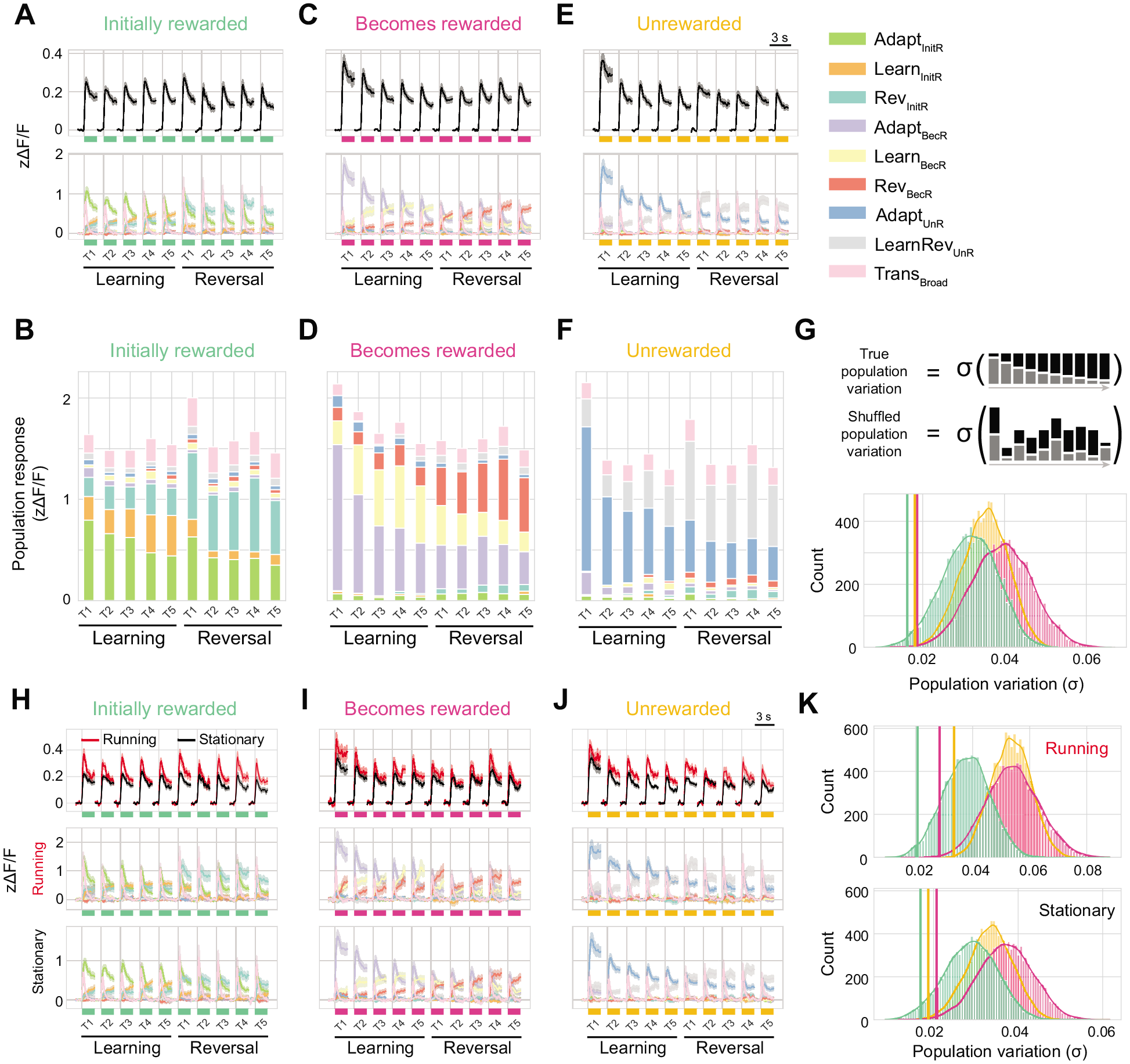
Average network activity is balanced across reward and locomotor contexts. A. Population response to the initially rewarded cue for each stage of learning, averaged across all neurons (top; cf. Fig. 2D) and across all neurons in a given component cluster (bottom). Average population responses were remarkably similar across stages, despite dominant underlying contributions from different groups of cells with distinct response dynamics within trials and across stages of learning. Note that, for this analysis, z-scored cue responses of each cell were not normalized to the peak response across stages. B. Stacked bar plot of population response magnitudes to the initially rewarded cue for each component cluster of neurons across each stage (compare to response magnitudes in A, bottom panel). The total activity across component clusters is stable across stages despite differences in which clusters were strongly responsive at each stage. C. Same as A, but for the cue that becomes rewarded after reversal. D. Same as B, but for the cue that becomes rewarded after reversal. E. Same as A, but for the cue that remains unrewarded. F. Same as B, but for the cue that remains unrewarded. G. Across-stage population average response stability was greater than expected by chance. Bottom: standard deviation of population response averages across stages was calculated before (vertical lines) and after shuffling cells by stage (distributions, n = 10,000 shuffles). Cells belonging to the same component-cluster were shuffled as a group (top). The observed variation across stages was significantly less than in the shuffled data, indicating that cue responses of cells in different component clusters balance one another for any given stage. Initially rewarded cue: p = 0.0115; Becomes rewarded cue: p = 0.0015; Unrewarded cue: p = 0.0012; bootstrap permutation test. H. Top: same as top panel of A but plotted separately for averages across running (red) and stationary (black) trials. Middle and bottom: same as bottom panel of A but plotted separately for averages across running trials (middle) and stationary trials (bottom). I. Same as H, but for the cue that becomes rewarded after reversal. J. Same as H, but for the cue that remains unrewarded. K. Standard deviation of population mean across stages was calculated as in G for averages across running trials (top) and stationary trials (bottom). The balanced population responses to each cue across stages was greater than expected by chance when considering running and stationary trials separately. Running: Initially rewarded cue: p = 0.0187; Becomes rewarded cue: p = 0.0024; Unrewarded cue: p = 0.0008. Stationary: Initially rewarded cue: p = 0.0312; Becomes rewarded cue: p = 0.0051; Unrewarded cue: p = 0.0033; bootstrap permutation test. See also Figure S5.

We also observed similar stability in average population responses to the other two cues, with the exception of the first stage of learning, for which net population responses were somewhat stronger. This increased response to these two cues (Fig. 5C-F) but not to the initially rewarded cue (Fig. 5A-B) could be explained by the additional pre-training sessions prior to the first stage of training on the three-cue task, in which only the initially rewarded cue was presented (together with unconditional delivery of reward; Fig. S5D-E). This analysis further showed that adapting cells in particular show an exponential decrease in response magnitude that continued across thousands of cue presentations across sessions (Fig. S5E).

We quantified whether the stability of net population activity due to this balancing between different clusters of cells that were responsive at different stages of learning exceeded the balancing expected by simply averaging many neurons together. For cells assigned to each component, we randomly shuffled the response time courses across stages (Fig. 5G, top). We found that the net response across all cells was indeed less variable across stages than for the shuffled data (Fig. 5G, bottom).

This same network-wide balance in average population response across learning stages was also present when separately considering trials involving a running or a stationary locomotor context (Fig. 5H-K), despite largely distinct sets of responsive neurons in these two contexts (Fig. 4). Overall, net responses were similar across these two conditions, albeit somewhat stronger during running (Fig. 5H-J), consistent with balanced sets of neurons predominantly driven by cues during running or stationary states (Fig. 4C-D, Fig. S5B). Taken together, these results suggest tight homeostatic regulation of overall network responsivity to cues in POR across stages of learning and reward and locomotion contexts, despite a division of labor where distinct neurons encode cues in different conjunctions of contexts.

### Diverse cue responses to stimulus offset across stages of learning and relearning

We also observed excitatory cells that were activated with diverse within-trial and across-stage dynamics during the period *following* stimulus offset (e.g., Fig 6A). Offset responses are understudied in the visual system, but recent evidence suggests that offset responses in sensory cortex may reflect mismatch between sensory predictions and sensory inputs (Hertäg and Sprekeler, 2020; Jordan and Keller, 2020; Keller and Mrsic-Flogel, 2018). Offset responses might simply signal any unpredicted transition in the visual environment (e.g., a stimulus ending). Alternatively, such responses could be selective to the offset of a specific cue in a particular reward context, similar to onset cells. We therefore examined in greater detail the sensory properties of these cells across reward contexts. We observed a surprisingly large number of neurons (445 out of 7824 neurons that were tracked across multiple days from 7 mice, see Methods) that were significantly and preferentially activated following the offset of the stimulus (Fig. 6A-B; using conservative criteria for responsivity), as compared to the 1799 cells driven preferentially during stimulus presentation (Figs. 1-2). When we sorted offset cells by their preferred cue and by their peak response latency following cue offset, we observed cells that were either broadly or sharply tuned, and exhibited a range of response shapes from fast and transient to slower and more sustained (Fig. 6A-B). This functionally diverse set of offset cells represents a sizeable proportion of all cells exhibiting trial-modulated activity.

**Figure 6.**
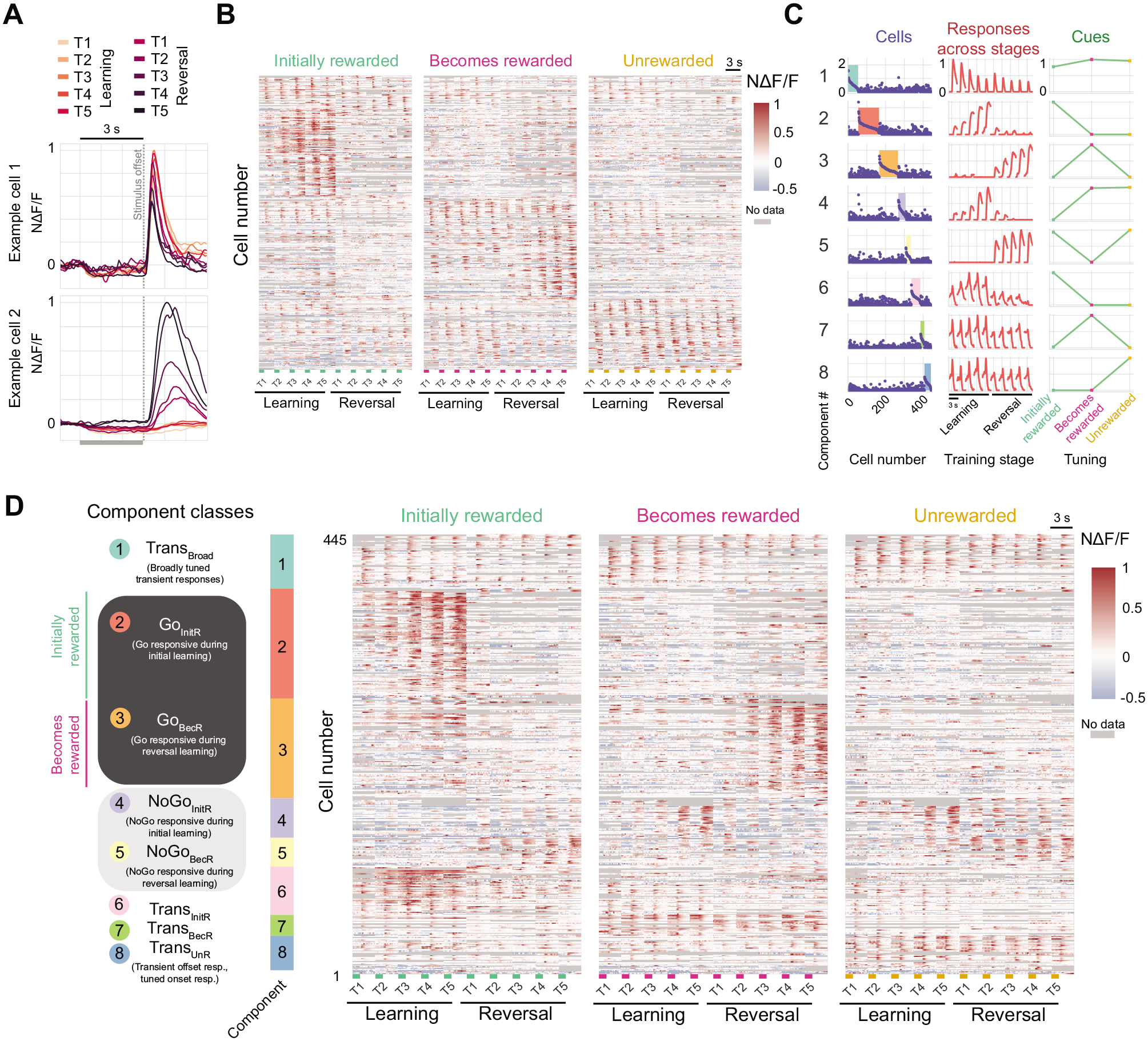
Evolution of responses in distinct classes of offset neurons across learning and relearning. A. Example traces from two stimulus offset-responsive neurons showing changes in responses across stages of learning. B. Heatmap of mean trial responses for each of 10 stages of learning for cells driven at stimulus offset (n = 445). Responses are normalized to the maximum response across stages and cues for each cell. Cells are sorted first by the cue that evoked the largest magnitude response, and then by their peak response latency from stimulus offset. Gray bars at bottom indicate the last second of visual stimulus before the offset. C. A tensor component analysis model with eight components captures distinct within-trial and across-stage response dynamics both with clear increases or decreases in response magnitude following reversal in a subset of components (see also Fig. S6). Plotting conventions are same as in Fig. 2C. D. Offset cells were classified as Go responsive (dark gray shading), NoGo responsive (light gray shading), or as transient (upper cluster and lowest 3 clusters). While all transient component clusters contained broadly tuned offset responses, the lowest three clusters in D also show ramping responses to one of the three cues. Offset cells were clustered according to maximum cell factor weight across all eight components (see shaded, colorbars in left column of C and corresponding component numbering at left of D). Cells were first sorted by cue preference and then by cell factor weight. n = 445 cells pooled across n = 7 mice. Gray bar indicates the last second before stimulus offset. See also Figure S6.

As with onset cells, the within-trial and across-stage dynamics in the population of offset cells were typically well described using a tensor component analysis model with eight components (Fig. 6C-D; the general classes of offset components were qualitatively similar when models with ranks greater than or less than eight were considered, Fig. S6). As with onset cells, we classified offset cells into functional categories by assigning each cell to the component that best described that cell’s activity (shaded regions in Fig. 6D, left column), and resorted the cell-by-cell heatmap in Fig. 6B by grouping cells assigned to each of the eight clusters (Fig. 6D). This showed that each of the offset components was faithfully reflected by the activity of many individual cells.

We observed four classes of offset components. First, we observed a component with broad tuning across the three cues and rapid, transient offset responses that were fairly consistent across all stages of learning and relearning (“Trans_Broad_” component). Second, we observed components that showed longer-lasting responses following the offset of the initially rewarded “Go” cue during high-performance stages of learning (“Go_InitR_” component), or following the offset of the cue that became rewarded following reversal learning (“Go_BecR_” component), indicated by dark gray shading on the left of Fig. 6D. Roughly half of all offset cells were associated with one of these two Go components. These components and associated cells were not driven early in learning, but showed robust responses during high-performance stages of learning — stages where task predictions and expectation of reward have developed. Critically, these cells were not directly driven by lick-related motor behavior or reward delivery, as they typically responded following a rewarded cue before or after reversal, but not both. Third, we observed components that responded following offset of both cues that did *not* predict rewards, reflecting a category-selective pooling of responses to the two “NoGo” cues (indicated by light gray shading in Fig. 6D, left). One component predominantly responded prior to reversal, after offset of the two NoGo cues that were not initially rewarded (“NoGo_InitR_” component). The other predominantly responded following reversal, after offset of the two NoGo cues that did not become rewarded following reversal (“NoGo_BecR_” component). Fourth, we observed three other components, each of which had a broadly tuned, fast transient offset response (as with the Trans_Broad_ component), but also exhibited a tuned, slow ramping onset response upon presentation of one of the three cues.

We quantified the above differences between clusters of offset neurons at the mouse-by-mouse level by averaging cells assigned to each cluster, using similar analyses as applied to onset neurons in Fig. 3. On average, we observed similar responses in each mouse from the cluster of offset cells assigned to each offset component (Fig. 7A), suggesting that these classes of offset cells were consistent across mice. We found that the clusters of cells with broadly tuned and transient offset responses all exhibited shorter onset latency with steeper slopes and were more transient than for Go or NoGo clusters of offset cells (Fig. 7B-D). Cell clusters also showed a marked difference in the sustainedness of their offset response (Fig. 7D, S7A). These cell clusters also differed in the learning stage where peak offset responses were observed (Fig. 7B, E), as is evident from examination of individual cells (Fig. 6E). As with onset cell clusters, we found that offset cell clusters with shorter response latency and faster slope (i.e., broadly tuned, transient offset neurons) showed less modulation by the operant response of the mouse (lick or no-lick) than the Go and NoGo clusters across trials (Fig. 7F-H). These findings show that “Go” offset cells with stronger offset responses when a cue was associated with reward are particularly strongly driven on correct trials during high performance stages (Fig. 7E,G-H, Fig. S7B) — precisely when the animal is likely to strongly expect a salient outcome. Similarly, “NoGo” offset cells with stronger offset responses when a cue was not associated with reward are particularly strongly driven on correct trials during high performance stages (Fig. 7E,G-H, Fig. S7B) — precisely when the animal likely expects that withholding licking will prevent a salient outcome (such as quinine delivery).

**Figure 7.**
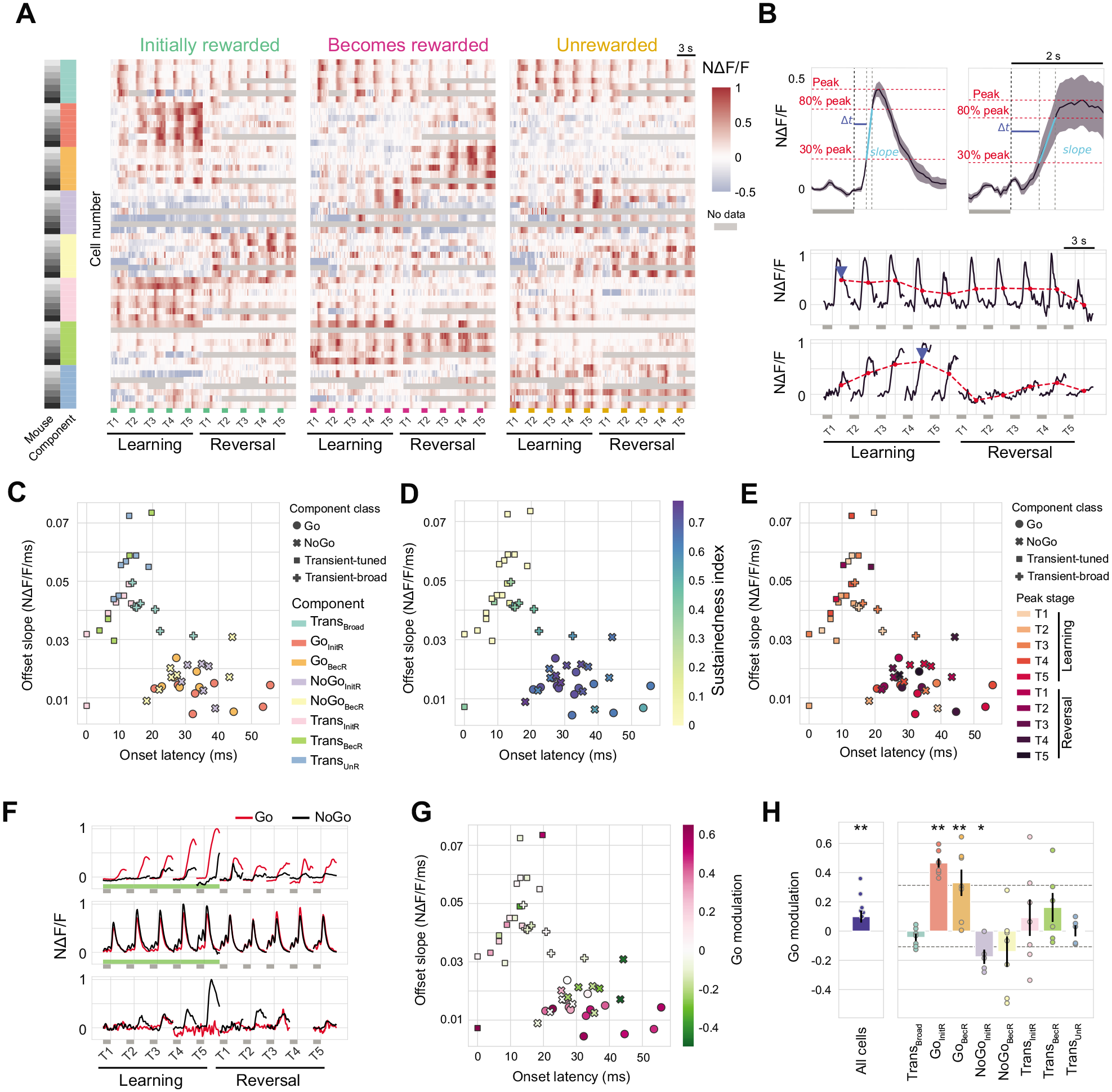
Offset responsive neurons with distinct within-trial and across-learning dynamics show differential sensitivity to reward context. A. Average response across cells from each mouse that belonged to each component cluster. Each mouse yielded qualitatively similar cells belonging to each component cluster. Results are sorted as in Fig. 6D. Gray bar indicates the last second of visual stimulus before the offset. B. Top: estimation of offset latency and offset slope for two example response traces (averaged across stages). Offset latency (blue line) was calculated as the first crossing of a threshold (30% of peak response) following stimulus offset. Offset slope (cyan line) was calculated between 30% and 80% threshold crossings (red lines). Bottom: estimation of mean offset response per stage (red line) and of maximally responsive stage (i.e., “Peak” stage; blue arrowhead). C. Scatter of onset latency versus onset slope. Each data point represents one mouse and component cluster. The shape of each data point denotes whether the component was considered a “Go,” “NoGo,” “Transient-tuned,” or “Transient-broad” component. Color denotes component cluster. D. Same as C, except that color indicates the “sustainedness” index of the average stimulus response of that component cluster. E. Same as C, except that color indicates the peak stage of the response across learning. F. Response to the initially rewarded cue for three example cells, averaged across all Go trials (red) or NoGo trials (black) in each stage of learning. Green bar indicates period where visual stimulus is rewarded. Grey bar indicates the last second of the visual stimulus. G. Same as C, but color indicates the average Go modulation of cells in each component cluster (Go modulation = [Mean Go resp. - Mean NoGo resp.]). H. Difference in normalized response magnitude for Go trials versus NoGo trials for each component cluster. Error bars: SEM across mice. **p < 0.001, *p < 0.05; bootstrap test over Go/NoGo trial averages, Bonferroni corrected for components (including “All cells”). Dashed line: 95% confidence interval estimated using bootstrapping across components. Error bars: SEM across mice. See also Figure S7.

We observed additional similarities with the findings for onset neurons. First, Go and NoGo offset neuron responses were quite stable across neighboring stages of high performance, when task rules were stable (Fig. S7C, top). In contrast, Go and NoGo offset neurons showed large changes in response magnitude in neighboring stages *across* reversal, yet this was not the case for the clusters of broadly tuned, fast transient offset neurons (Fig. S7B, bottom). Taken together, these findings show that the diverse clusters of offset neurons also divide into two main types: neurons that respond at cue offset regardless of the meaning of the cue, and neurons that respond at cue offset depending on the current reward context (i.e., whether a given cue is currently associated with reward or not). These two types of offset responses may reflect different kinds of negative predictions errors regarding the unexpected disappearance of a cue that is either visually salient regardless of task context, or that has acquired motivational salience in a particular task context (see Discussion and Fig. S7C). These data support the notion that POR neurons encode cue information in both a context-independent and context-dependent manner, not just during stimulus presentation but also after stimulus offset.

## DISCUSSION

Here, we have used chronic calcium imaging to track excitatory neurons in layer 2/3 of POR throughout all stages of cue-outcome learning and reversal learning. Some POR neurons exhibited responses to specific cues that are independent of reward context. These short-latency responses were present even prior to association of any cues with rewards, and likely reflect the bottom-up salience of the stimuli, consistent with their progressive response adaptation across thousands of trials. Meanwhile, other neurons exhibited longer-latency, sustained responses to specific cues in particular conjunctions of contexts (locomotion/stationary, rewarded/unrewarded). Below, we consider our findings in the context of previous studies of visual processing and of potential contributions of inputs from other brain regions to contextualization of sensory information in POR. We then discuss technical advantages and limitations of our experimental approach to imaging and analyzing data from the same neurons across many weeks.

### Conjunctive contexts for cue learning

We found that specific sets of POR excitatory neurons exhibit delayed-onset, sustained cue responses limited to specific conjunctions of reward and locomotor contexts. The number of neurons driven in each conjunction of contexts (e.g., cue is rewarded or non-rewarded; mouse is running or stationary) was generally balanced. This resulted in a similar total cue-evoked increase in activity across all recorded neurons in each conjunction of contexts, consistent with previous demonstrations of similar network homeostasis in other regions such as the hippocampus (Buzsáki et al., 2002; Slomowitz et al., 2015) and primary visual cortex (Benucci et al., 2013). This balance was driven in part by gradual recruitment of subsets of neurons that responded to a cue as it became unrewarded, in addition to recruitment of other neurons driven by a newly rewarded cue. Neurons driven selectively by a cue in contexts where the cue has become both familiar and non-salient may constitute a kind of extinction trace, as observed in other brain areas (Mount et al., 2021; Reinert et al., 2021). Also surprising was the substantial set of excitatory POR neurons that responded to cues only when the mouse was stationary, in contrast to the strong bias in responses during locomotion observed in V1 (Andermann et al., 2011; Erisken et al., 2014; Niell and Stryker, 2010; Pakan et al., 2016; Saleem et al., 2013). Our findings of learned cue responses that depend on conjunctions of contexts may extend our understanding of contextual processing of cues in visual and association cortex, which previously focused largely on spatial context (Bucci et al., 2000; Furtak et al., 2012; Norman and Eacott, 2005; Saleem et al., 2018). Human fMRI and MEG studies have also demonstrated selective encoding of objects in specific *task* contexts in lateral visual cortex, but not in early visual cortex (Harel et al., 2014; Hebart et al., 2018). Notably, the late onset of contextual influences in these studies is consistent with our finding that context-dependent cue responses exhibit slower onset latencies.

The observed selectivity of many POR neurons for conjunctions of locomotion and reward contexts begs the question of whether locomotion is truly an internal context in a similar sense as, for example, spatial context. As in the majority of V1 neurons (Niell and Stryker, 2010), locomotion-dependent enhancement of cue responses in POR may be explained by increases in cortical arousal and neuromodulatory inputs to cortex (Larsen et al., 2018). However, increased arousal would not explain the substantial set of excitatory neurons in POR that are selectively driven by visual cues in stationary contexts. Moreover, while increased responsivity in stationary contexts could be due to mismatch signals related to unexpected optic flow (Jordan and Keller, 2020; Keller et al., 2012; Roth et al., 2016; Zmarz and Keller, 2016), our observation that cue responses are additionally dependent on reward context is not easily explained solely in terms of sensory-motor mismatch errors. Thus, it may be useful to consider locomotion as a distinct internal context (Erisken et al., 2014) comprised of distinct tactile, proprioceptive, and interoceptive sensory signals — much as hunger and satiety may also be viewed as internal contexts (Burgess et al., 2018; Burgess et al., 2016; Livneh et al., 2017). These internal signals may complement the spatial signals that are traditionally viewed as important for contextual learning. Indeed, memory retrieval can often depend on whether the association is learned and retrieved in the same interoceptive context (Kennedy and Shapiro, 2004). Both self-motion and interoceptive signals may contribute to the finding that distinct sets of local and distal cortical regions are correlated with individual visual cortical neurons across stationary or locomotor states (Clancy et al., 2019). Learning-related association of cues with inputs from one of these two sets of neurons may provide a neural substrate for locomotor context-dependent learning in visual cortex. While the drivers of cue-in-context encoding in POR neurons remain unknown, the slow-onset, sustained nature of these responses suggest an important role of top-down feedback (Keller et al., 2020; Makino and Komiyama, 2015), including from the posterior parietal and retrosplenial cortices, the lateral amygdala, and the entorhinal cortex (Burgess et al., 2016; Burwell and Amaral, 1998; Meier et al., 2021; Wilson et al., 2013).

The emergence of selective encoding of cues in conjunctions of contexts by neurons with maximal responses during later, higher-performance stages of training is consistent with the orthogonalization of sensory response patterns during learning (Failor et al., 2021; Poort et al., 2015), and may allow for more efficient decoding of objects in specific contexts (versus a more distributed representation). The finding that clusters of context-related cells are driven by a rewarded cue or by a non-rewarded cue during initial learning, but not after reversal, may reflect a long-term memory of the initial cue-context association (Namboodiri et al., 2019). Such memories may be adaptive in situations where there is a return to the initial context (testable in future studies). This is in contrast to studies that have shown individual cells selective for rewarded stimuli regardless of reward context across reversal in orbitofrontal cortex (Banerjee et al., 2020) and POR (Ramesh et al., 2018). The current study provides a much richer description of learning-associated changes in network activity than was possible in our previous recordings of POR neurons across a small number of days following viral expression of GCaMP6f (Ramesh et al., 2018).

In addition to cells with context-dependent encoding of cues, we also observed cells that encoded specific visual stimuli regardless of context, and often even prior to association of the stimuli with salient outcomes (e.g., in naïve mice). These neurons were adapting both in terms of their increasingly transient and increasingly smaller amplitude responses across stages, in a manner that was minimally affected by the change in cue contingencies. We speculate that these neurons may support a stable, veridical representation of object identity regardless of context, but with response magnitudes that reflect gradual decreases in the unexpected bottom-up salience (e.g., familiarity) of these stimuli after thousands of stimulus presentations (Freedman et al., 2006; Li et al., 1993). This division of labor between context-dependent and context-independent neurons in POR could allow both stability in object identification and flexibility in guiding how to react to such objects in particular contexts.

### Do stimulus offset responses reflect prediction errors?

Interestingly, about 20% of cue-driven neurons in POR respond preferentially following stimulus offset. In most cases, these responses were not strongly related to motor behavior during this period. For instance, offset neurons were often responsive on non-rewarded trials that lacked motor behavior. This was true both of broadly-tuned, fast transient offset neurons and slower-onset NoGo neurons that showed generalized responses across multiple cues not associated with reward in a given context (Reinert et al., 2021). Further, offset neurons that responded on Go trials mainly did so in a specific cue-reward context prior to or following reversal. Such response properties of context-specific Go or NoGo responses also constitute additional kinds of context-dependent cue responses in POR.

While visual offset responses are rarely examined in detail, studies in anesthetized cat visual cortex (Ferster, 1988; Liang et al., 2008) have proposed that offset responses may relate to the psychophysical phenomenon of visual persistence (Liang et al., 2008). This notion that visual responses may relate to predictions of persistence of a salient sensory stimulus, or mismatches between such predictions and actual sensory input, has been developed in several theoretical studies involving predictive coding (e.g., Keller and Mrsic-Flogel, 2018; Rao and Ballard, 1999). For example, Keller and Mrsic-Flögel recently built on the predictive coding framework to suggest that distinct neurons in upper layers of a given cortical area encode either the difference between a sensory stimulus and a prediction of that stimulus (S-P; a “Type 1” prediction error), or the difference between the prediction of that stimulus and the actual sensory input (P-S; a “Type 2” prediction error). As proposed in Fig. S7C, the responses of various offset cells in our dataset may reflect diverse types of Type 2 prediction errors. For example, in the case of a prediction that a high-contrast visual stimulus will remain on the screen, this prediction would be broadly tuned and would persist for a short delay following stimulus offset. In this case, a P-S computation would result in a fast, transient, broadly-tuned offset response that should be independent of the stage of learning (cf. upper cells in Fig. 6E; cf. offset responses in V1 neurons in Jordan and Keller, 2020). In contrast, a learned prediction that a rewarded visual stimulus is associated with a prolonged, multisensory event would result in an offset response that is tuned to the rewarded cue and persists for seconds following visual cue offset (cf. Go cells in Fig. 6E). Indeed, lateral amygdala feedback axons in POR show biased responses to the rewarded cue, and these responses appear sustained for seconds following cue offset (Burgess et al., 2016). Similar notions regarding distinct predictions could give rise to other offset types that we observed. For example, the neurons that showed generalized NoGo offset responses after both unrewarded cues could be driven by feedback predictions from generalized NoGo onset neurons in prefrontal cortex (Reinert et al., 2021). Regardless of whether these neurons reflect prediction errors or not, they demonstrate that the encoding of cues in very specific contexts occurs not only in large subsets of onset-driven neurons but also in rarely examined, offset-driven neurons.

### Technical advantages and limitations

Initial learning and reversal learning of an operant visual Go/NoGo task typically requires thousands of trials across weeks of training in mice (e.g., Fig. S1B, Ramesh et al., 2018; Reinert et al., 2021). While challenging for the experimenter, this “slow-motion” learning trajectory can be advantageous. Specifically, it enables averaging of visual responses across many hundreds of trials at each stage of learning (Fig. 1), allowing robust estimation of the evolution of visual responses properties. This benefit comes at the cost of having to track the same neurons across many weeks, from prior to any associations with rewards/punishments all the way through learning and relearning. This was not possible in our previous reversal learning experiments (Ramesh et al., 2018), which did not investigate initial learning and which tracked neurons over a much smaller number of sessions due to the use of viral expression of calcium indicators with progressively higher expression levels across weeks. We overcame this challenge by recording from transgenic mice that stably expressed GCaMP6f (Madisen et al., 2015) across tens of sessions (median: 25 sessions/mouse).

We used activity-based algorithms to identify and segment neurons in each recording. Using this approach, a neuron’s mask was sometimes missing in a subset of sessions. While we cannot strictly rule out technical considerations such as slight differences in the tilt of the imaging plane across days, we are confident that in most cases, a cell remained in the field of view but was silent during these sessions (Video S1). Indeed, within a given learning stage, sessions in which a cell mask was missing tended to occur in close proximity to sessions in which the cell was present but mostly silent. The strategy of averaging neuron responses across sessions within a stage of learning (cf. Poort et al., 2015) allowed us to recover cells even during relatively quiet periods, which was critical to our findings that some neurons are only driven by cues in certain task contexts. Our ability to classify neurons despite occasional missing data from learning stages in which the neuron’s cell mask was entirely absent benefited from the use of tensor component analysis, which can handle missing data points (Williams et al., 2018).

In our study, our use of averaging across sessions within a stage of learning allowed us to demonstrate reliable cue encoding across stages (e.g., in adapting clusters of cells). While averaging across sessions is useful from an experimental standpoint (e.g., averaging over small errors in the sectioning of a field of view each day), it also can obscure potential daily or hourly biological fluctuations (e.g., in response gain). Thus, while recent studies have noted the presence of “representational drift” and across-day changes in responsivity of neurons encoding cues, actions, or locations (Deitch et al., 2020; Driscoll et al., 2017; Huber et al., 2012; Schoonover et al., 2021; Sheintuch et al., 2020), our data averaging multiple days of data per stage of learning suggest that many functional groups of excitatory neurons show stable, long-term sensory encoding. This was particularly true of cells that are insensitive to changes in reward context (e.g., adapting cells), but was also true of context-sensitive cells, once cues and associated reward/locomotor contexts had been stably learned (e.g., between T4 and T5 stages of initial learning). Previous studies have highlighted the utility of such stable representations for recall of specific memories (Kitamura et al., 2017; Liu et al., 2012; Yiu et al., 2014).

Far from minor technicalities, these considerations regarding analysis of data across tens of days of imaging are essential for accurately assessing the highly dynamic nature of sensory cortical responses across learning. More generally, our findings highlight the value of high-dimensional datasets that track responses in the same large set of neurons across the entire arc of learning for advancing our understanding of the full range of computations performed in association cortex.

---

**Figure S1.**
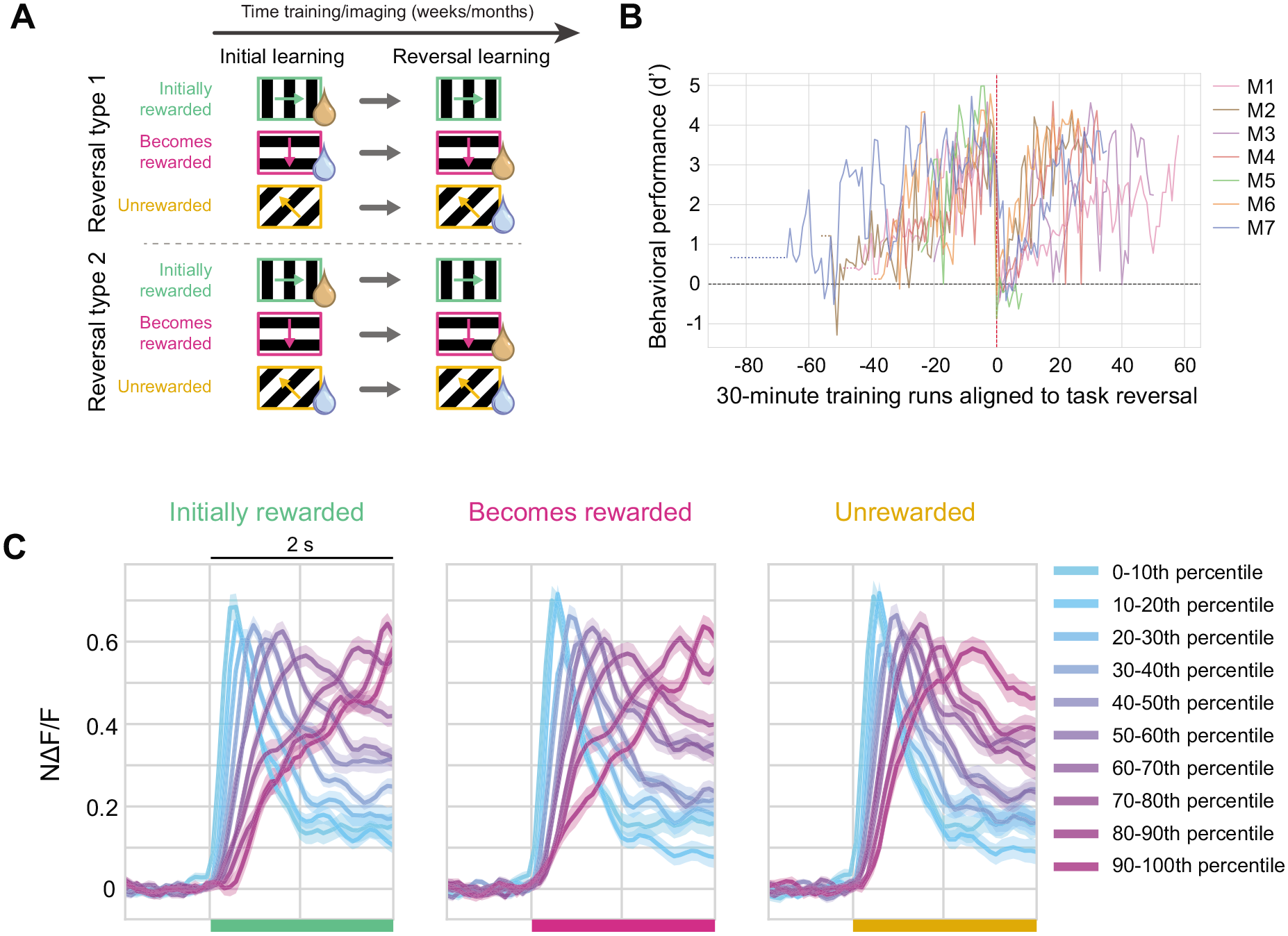
Behavioral performance across learning and reversal learning, related to Figure 1. A. Initial learning and acquisition of the task was followed by a change in cue-outcome contingencies during reversal learning. For each mouse, we used one of two types of reversal: either all cues changed across reversal (top), or the punished cue was held constant while the rewarded and neutral cues were swapped (bottom). See also Figure 1A-B. B. Behavioral performance for each mouse (“M1” to “M7”) was calculated across 30-minute imaging runs (1-6 runs per day). Dotted lines indicate passive viewing runs prior to onset of training (and thus lacking an estimate of performance, d’). C. Average cue-evoked response across subsets of neurons ranked by peak response latency (as in Fig. 1G). Neurons that prefer distinct cues show similar response time courses across a range of peak response latencies. Error bars: SEM across cells.

**Figure S2.**
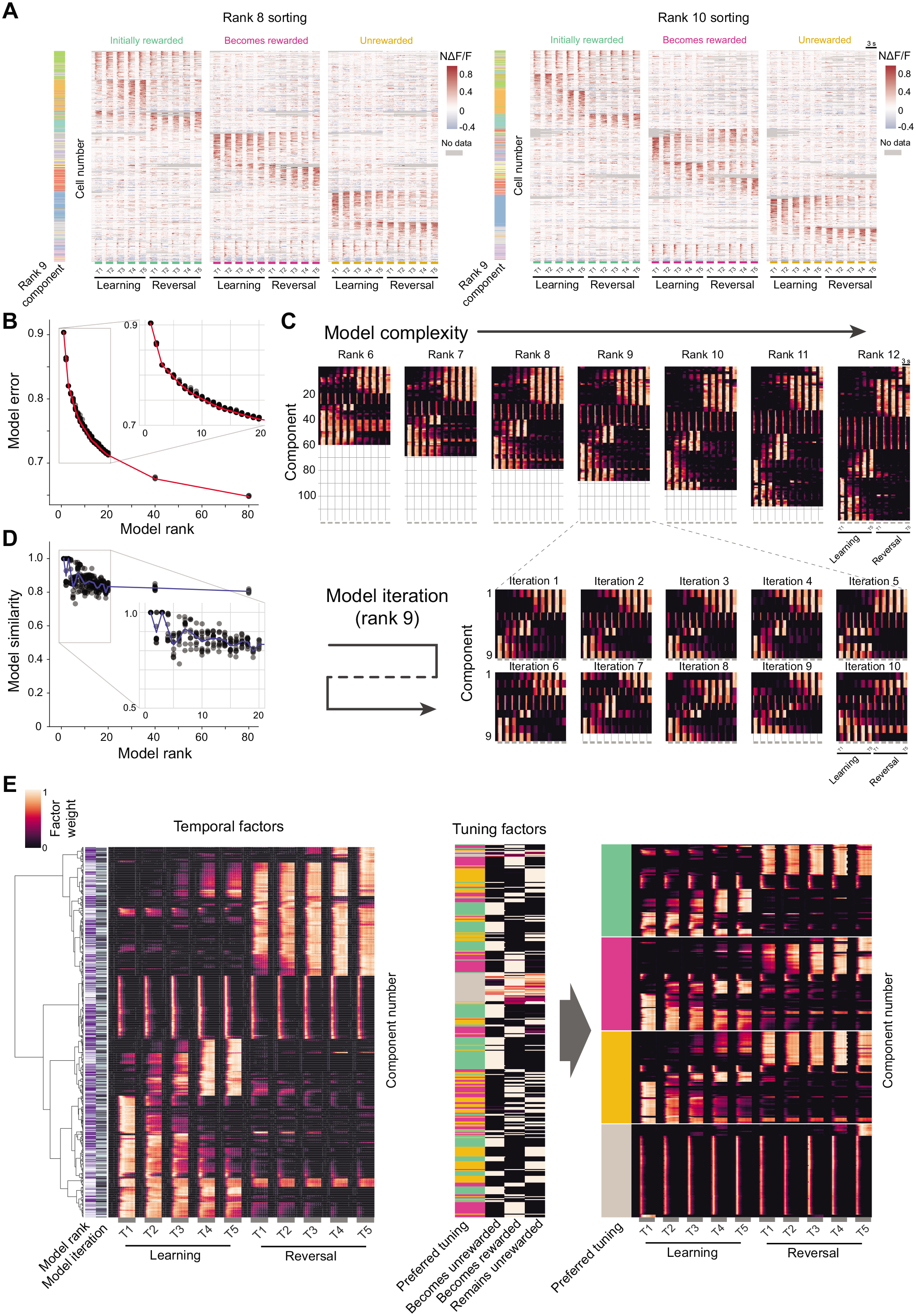
Tensor component analysis model selection and cell clustering, related to Figure 2. A. Example heatmaps as in Figure 2D, sorted by maximum cell factor weight for a rank 8 TCA model (left) and a rank 10 TCA model (right). Colorbars to the left of each heatmap indicate the classification of each cell based on the rank 9 model used in Figure 2D. Neurons are often assigned to a common group across different models. Thus, this analysis demonstrates qualitatively similar functional clustering of neurons when using rank-8, rank-9, or rank-10 TCA models. B. Normalized model reconstruction error across increasing model ranks of nonnegative TCA models. Ten optimization iterations (i.e., fits of the model with different random initializations, indicated by gray dots) of TCA were run for each rank, and the best of these was selected for subsequent analyses (red line). Models between rank 6 and 12 were considered to exhibit a reasonable range of performance without overfitting (i.e., a “knee”). C. Top: models of differing rank (10 optimization iterations per rank) show qualitatively similar components even as model complexity increases. Components are sorted using linkages from hierarchical clustering of temporal factors for all iterations and all sizes of models (using ranks of 6-12 components). Bottom: optimization iterations for rank-9 models ordered from lowest reconstruction error to highest reconstruction error. Independent iterations of randomly initialized TCA models produce qualitatively similar dynamics across sets of components. TCA can produce solutions that are simpler than the allotted model rank so it is possible to have fewer than 9 components (e.g., iterations 7 and 9). D. Model similarity as a function of model rank. Similarity, a measure of the distance between the optimal permutations of TCA factors for two models, for each optimization iteration is plotted compared to the lowest error model. Gray dots correspond to the same models as in B. Mean similarity across iterations is shown in blue. Similarity scores above 0.8 were considered qualitatively and quantitatively similar (see Methods; Williams et al., 2018). E. Agglomerative hierarchical clustering of temporal factors for all iterations (n = 10 iterations per rank) and all model sizes (ranks 6-12) shows components with transient or sustained activity within a stage of learning as well as distinct changes across initial learning and reversal (left panel). By combining this with cue tuning factors for each component (center panel), components were separated into 4 groups based on preferred tuning (right panel). From these groupings, 9-10 clusters of components were categorized using hierarchical clustering. The broadly tuned component group (gray) contained largely fast, transient responses. For tuned component groups, there were typically 2-3 functionally distinct types of temporal factors per cue that were evident across a large number of models. This indicates that, for each cue, there were typically three classes of cells whose responses evolved in different ways across learning and reversal. These could be broadly categorized as those whose responses 1) are strongest early in initial learning and adapt over stages, 2) increase in magnitude during initial learning, or 3) increase in magnitude during reversal learning. Fewer additional components capture other diverse activity patterns across learning. We therefore concluded that a model rank of nine was a reasonable balance of qualitative activity patterns and the total component number in each group. Other ranks and iterations of TCA produced very similar results.

**Figure S3.**
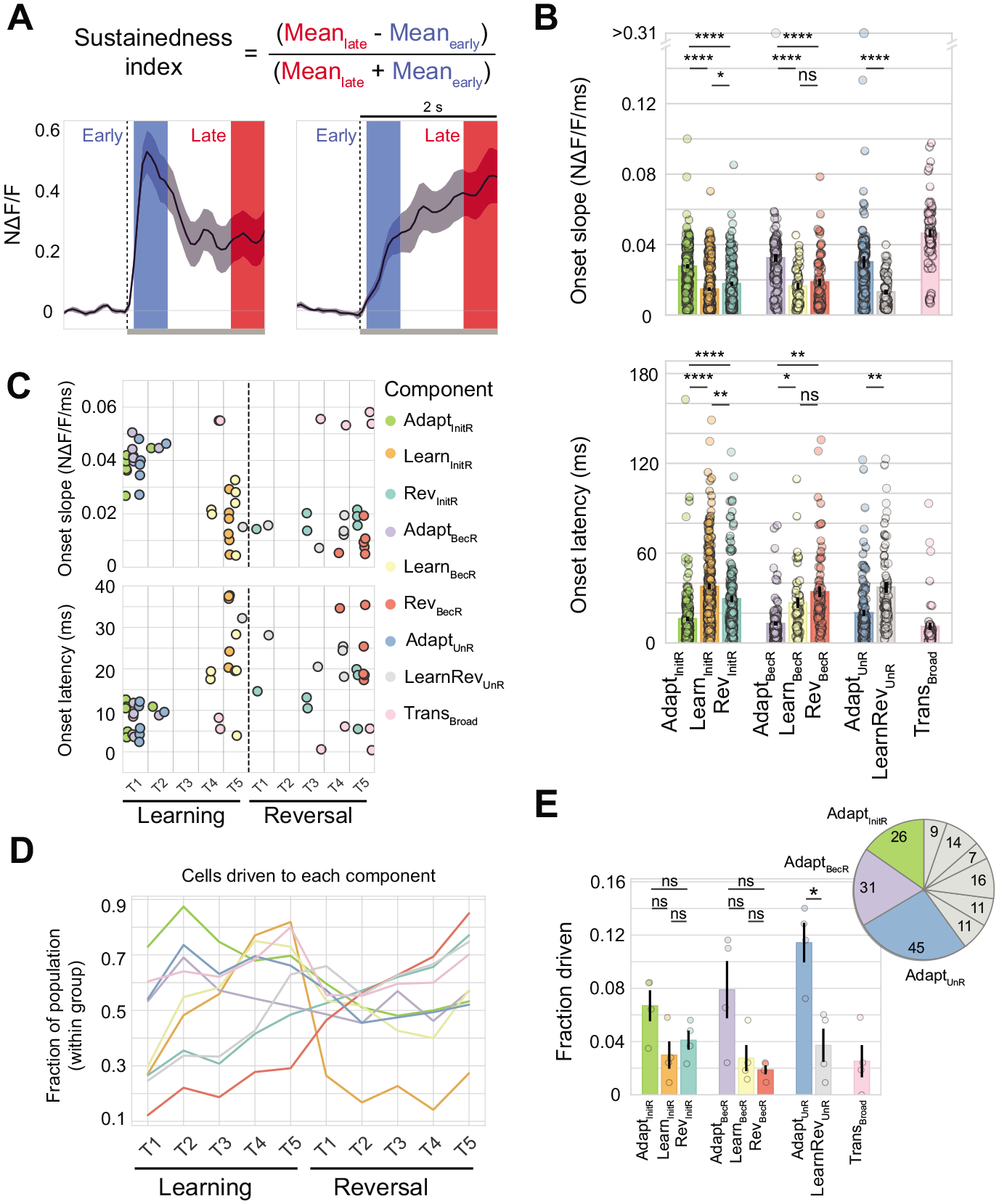
Additional analyses of onset-driven neurons across component clusters and mice, related to Figure 3. A. Calculation of sustainedness index for two example response traces (averaged across stages). This was calculated using the first 500 ms windows (at a 65 ms delay) of the stimulus presentation (“early”) and the last 500 ms of the 2 s stimulus window (“late”). Sustainedness index = [Mean_late_ -Mean_early_] / [Mean_late_ + Mean_early_]. B. Onset slope (top) and onset latency (bottom) summarized across all cells (n = 1799) according to component cluster (i.e., according to the functional cluster to which they were assigned. Comparisons were made between clusters of cells that share the same cue preference (*p < 0.05, **p < 0.005, ****p < 0.00005; two-sided Mann Whitney U test; Bonferroni corrected for comparisons within cue type). Error bars: SEM across cells. C. Onset slope (top) and onset latency (bottom) versus peak stage for mean responses for each component across all 7 mice (data are same as in Figure 3E). D. Proportion of cells within each component cluster that are driven at each stage of learning. Fraction of cells in each component cluster that were driven during cue presentation in naive mice, prior to conditioning (*p < 0.05; two-sided Mann Whitney U test; Bonferroni corrected for comparisons within cue type; n = 4 mice; error bars, SEM over cells) (left). Inset: Pie chart of numbers of driven cells belonging to each component cluster. Of the total number of cells driven naïve (n = 179 cells), cells from adapting component clusters made up more than half (102/179 cells). A majority of neurons driven in naïve sessions are from of the three adapting component clusters.

**Figure S4.**
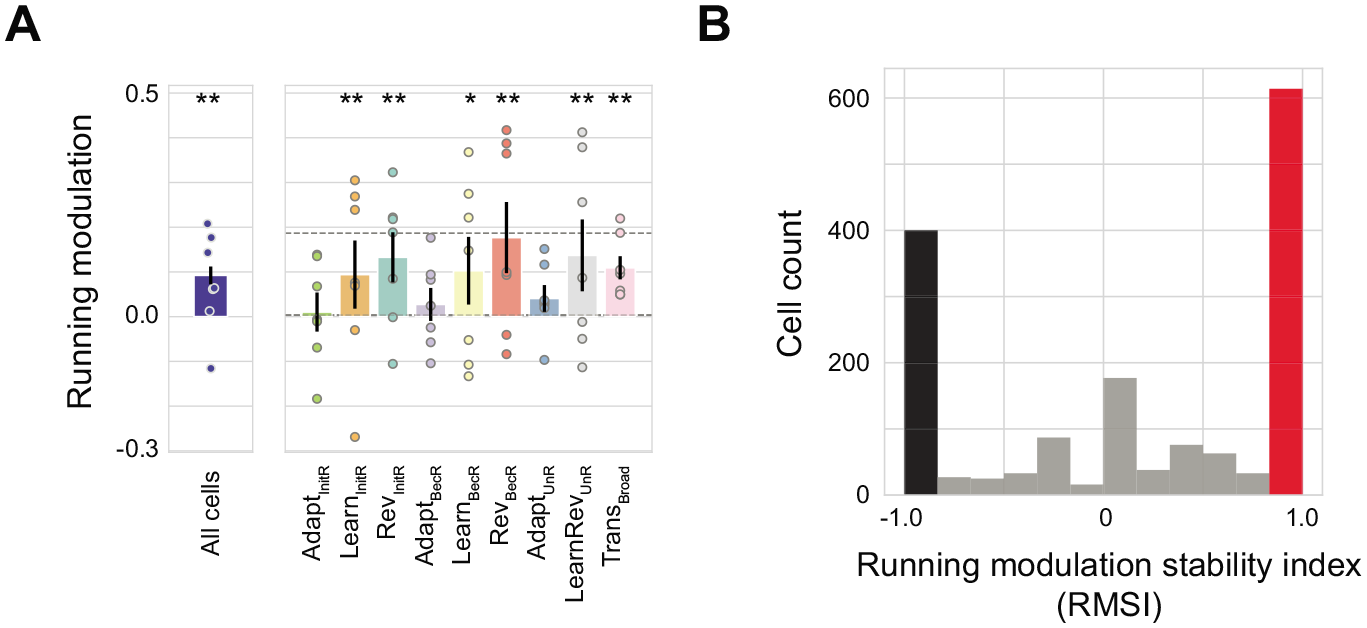
Characterization of modulation of cue responses by locomotion, related to Figure 4. A. Plot of average running modulation for each component cluster in each mouse, estimated as the mean response to all running trials minus the mean response to stationary trials, averaged across all cells within each component cluster. There was a small but significant net bias in response modulation by locomotion in adapting component clusters, and small positive biases in other clusters. Average running modulation was also calculated after averaging responses across all cells from all component clusters in each mouse (left, blue). *p < 0.05, **p < 0.001; bootstrap test over running/stationary trial averages, Bonferroni corrected for components (including “All cells”). Dashed line: 95% confidence interval estimated using bootstrapping across components. Error bars: SEM across mice. B. Running modulation stability index (RMSI) was calculated as RMSI = [(number of stages in which cue response during running trials was greater than during stationary trials) - (number of stages in which cue response during stationary trials was greater than during running trials)] / (total number of stages where the cell was driven by the cue). Cells with an RMSI of −1 consistently preferred stationary trials, while cells with an RMSI of 1 consistently preferred running trials across driven stages of training. Responses were estimated using the cell’s preferred cue.

**Figure S5.**
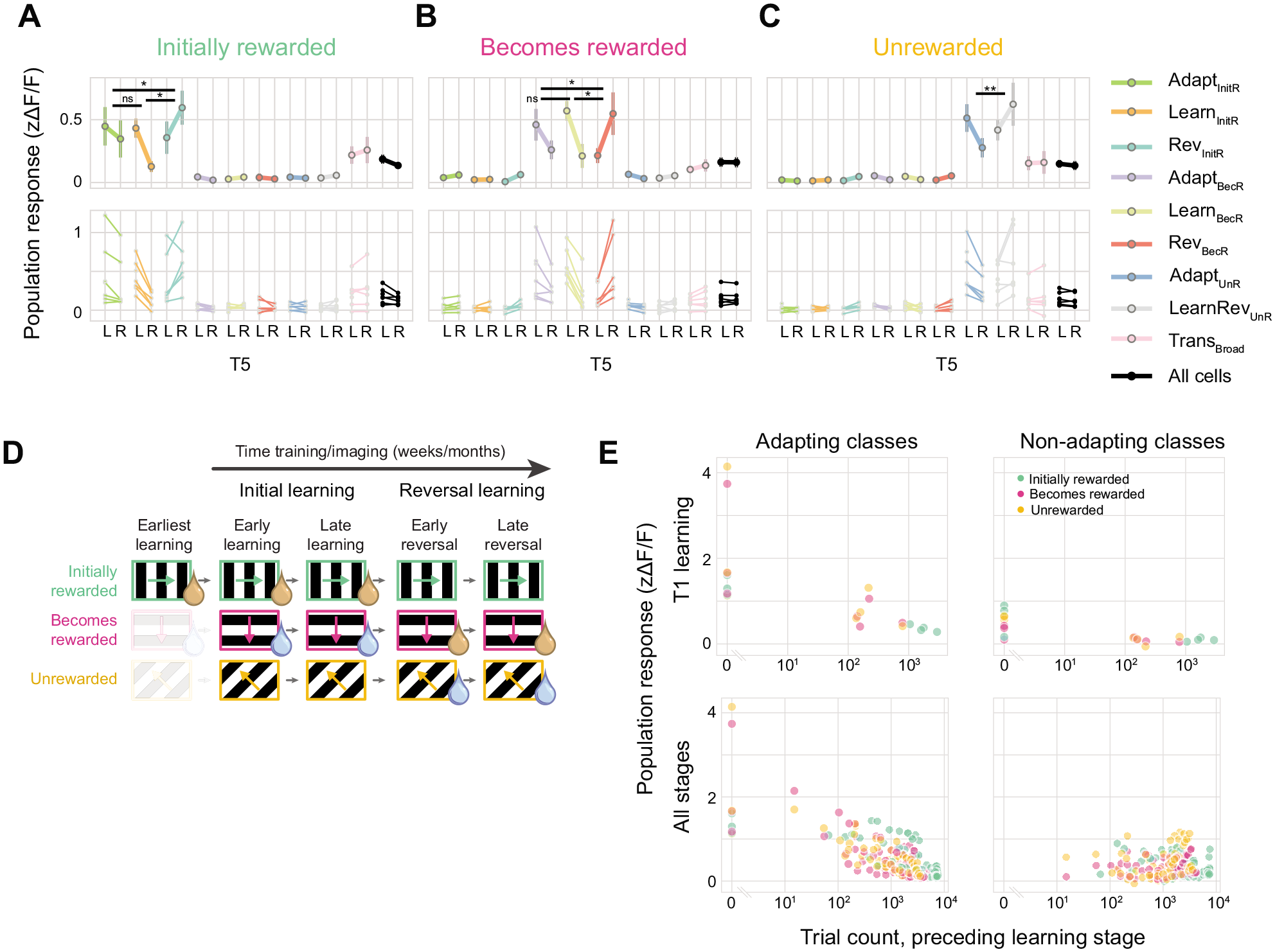
Balanced changes in component cluster responses across learning and adaptation effects of additional early training using a single cue, related to Figure 5. A. Magnitude of average response to the initially rewarded cue across cells in each component cluster during T5 learning (T5-L) or T5 reversal (T5-R), averaged across mice (top) or shown for individual mice (bottom). The overall mean population response across all cells (black) is stable across these distant stages of learning, even though distinct populations of neurons contribute to this signal during T5 learning versus T5 reversal stages. *p < 0.05, **p < 0.005; two-sided Mann Whitney U test for differences in T5 learning versus T5 reversal in each component cluster, Bonferroni corrected for multiple comparisons for each cue type. Error bars: SEM across mice (top). B. Same as A, but for the cue that becomes rewarded after reversal. C. Same as A, but for the cue that remains unrewarded. D. During the earliest stages of training (i.e., prior to training sessions included in other analyses), a large number of trials in which the initially rewarded cue was paired with unconditional reward (Pavlovian trials) or rewarded conditional on operant lick responses were used to incentivize initial performance on the task. During these initial sessions, the other two cues were not shown in order to sustain a high reward rate and minimize frustration during initial motor learning and cue-reward association. As described in Fig. 3F, we also presented equal numbers of trials of each of the three stimuli in the absence of reward or punishment outcomes in naïve sessions in 4/7 mice. E. Average response in each training stage versus the number of total trial exposures of each cue at the beginning of that stage. The average response magnitude across cells in each adapting component cluster at stage T1 of initial learning decayed exponentially across trial exposures (top left; one data point per mouse and adapting component), accounting for the differences seen in Figure 5A-F across stages between the consistently flat responses to the initially rewarded cue (green) and the additional adaptation observed for the other two cues (orange and pink). Other non-adapting component clusters did not show the same decay with trial exposure (top right). This same decay with cue exposure was apparent for adapting but not for non-adapting component clusters when considered across all stages of training (bottom; one data point per mouse, component cluster, and stage of learning).

**Figure S6.**
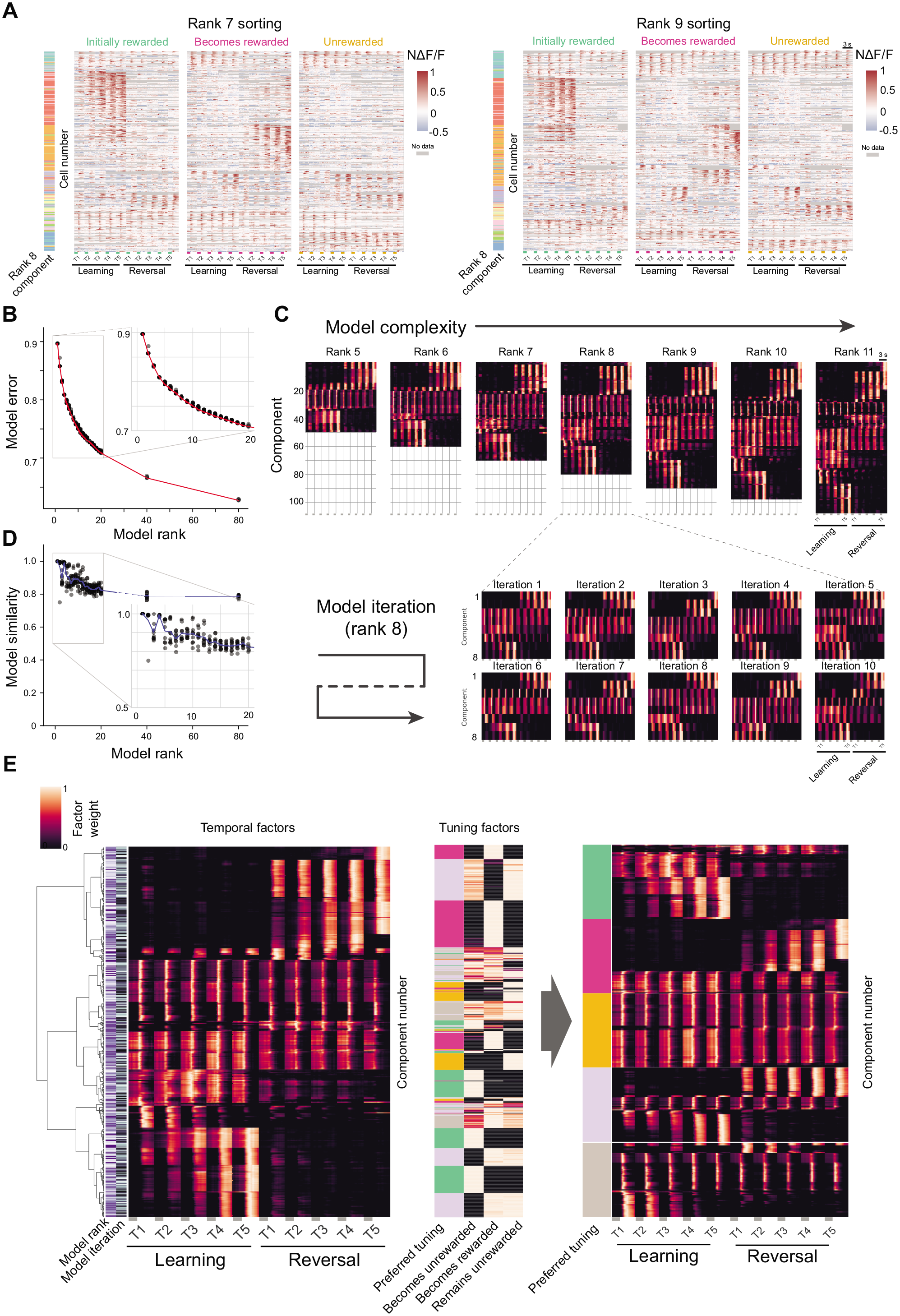
TCA model selection and clustering for offset cells, related to Figure 6. A. Example heatmaps as in Figure 6C, sorted by maximum cell factor weight for rank-7 (left) and rank-9 (right) TCA models. Colorbars to the left of each heatmap refer to the component cluster to which each cell belongs, as defined for the rank-8 model in Fig. 6C. Sorting reveals qualitatively similar clusters of neurons across these models of differing rank. B. Normalized model reconstruction error across increasing model ranks of nonnegative TCA models. Ten optimization iterations (i.e., fits of the model with different random initializations) of TCA were run for each rank (gray dots), and the best of these was selected for subsequent analyses (red line). Models between rank 5 and 11 were considered to exhibit a reasonable range of performance without overfitting (i.e., a “knee”). C. Top: models of differing rank (10 optimization iterations per rank) show qualitatively similar components even as model complexity increases. Components are sorted using linkages from hierarchical clustering of temporal factors for all iterations and all sizes of models (using ranks of 5-11 components). Bottom: optimization iterations for rank-8 models ordered from lowest reconstruction error to highest reconstruction error. Independent iterations of randomly initialized TCA models produce qualitatively similar dynamics across sets of components. D. Model similarity is plotted as a function of model rank. Similarity, a measure of the distance between the optimal permutations of TCA factors for two models, is plotted for each optimization iteration as compared to the lowest error model. Gray dots correspond to the same models as in B. Mean similarity across iterations is shown in blue. Similarity scores above 0.8 were considered qualitatively and quantitatively similar (see Methods; Williams et al., 2018). E. Agglomerative hierarchical clustering of temporal factors for all iterations (n = 10 iterations per rank) and all model sizes (ranks 5-11) shows components with transient or sustained activity within a stage of learning as well as distinct changes across initial learning and reversal (left panel). By combining this with tuning factors for each component (center panel), components were separated into 5 groups based on preferred tuning (right panel). From these groupings 8-10 clusters of components were qualitatively categorized. For tuned component groups there were two dominant activity patterns per cue with a large number of components across models. These could be broadly categorized as 1) responding to the stimulus and having a sharp offset response 2) responding during initial learning or reversal learning preferentially. While all three cues contained category-1 components, category-2 components only existed for cues that were initially rewarded or became rewarded. The broadly tuned component group (gray) contained largely fast, transient responses or, when their tuning was biased towards only two cues, delayed responses. Joint-tuned components (lavender) contained mostly delayed responses that transitioned across learning and relearning. We determined that a model rank of 8 was a reasonable balance of qualitative activity patterns and the total component number in each group. Other ranks and iterations of TCA produced very similar results.

**Figure S7.**
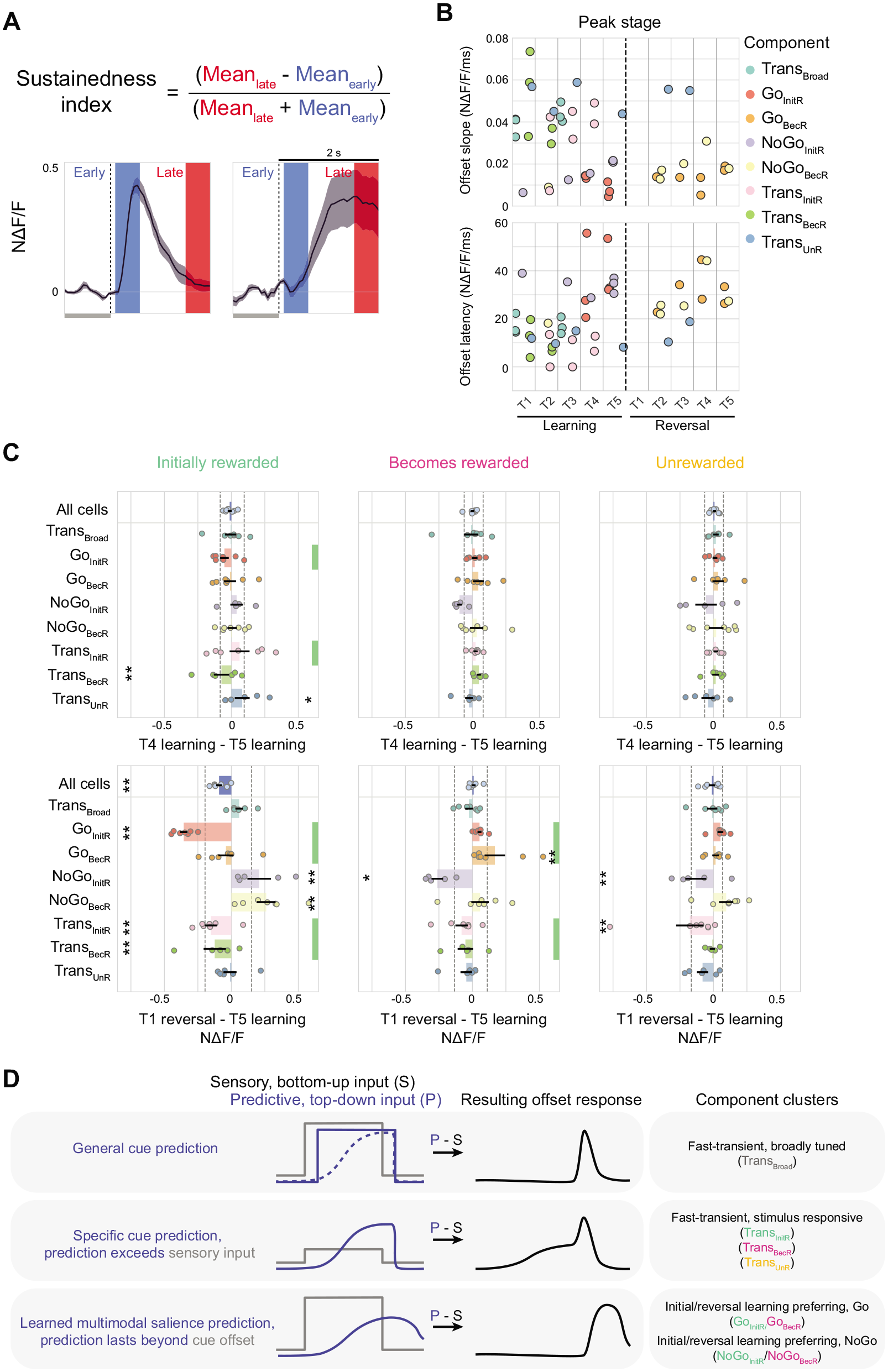
Additional analyses of offset responses and predictive coding framework, related to Figure 7. A. Calculation of sustainedness index for two example offset response traces (averaged across stages). This was calculated using the first 500 ms windows (at a 65 ms delay) after the stimulus offset (“early”) and the last 500 ms of the 2 s response window (“late”). Sustainedness index = [Mean_late_ - Mean_early_] / [Mean_late_ + Mean_early_]. B. Offset response slope (top) and offset latency (bottom) versus peak stage for mean response across cells in each component cluster and for each of the 7 mice (data are same as in Figure 7E). C. Difference in normalized response magnitude for average traces of each offset component between two neighboring stages of training that either straddled the reversal (top; T5 learning versus T1 reversal) or that both preceded the reversal (bottom; T4 learning versus T5 learning). Green bars indicate component clusters whose preferred cue is rewarded during at least one of the two stages. *p < 0.05, **p < 0.001; bootstrap test over stages, Bonferroni corrected for components (including “All cells”). Dashed line: 95% confidence interval estimated using bootstrapping across components. Error bars: SEM across mice. D. Future experiments might consider the predictive coding framework to explain the diversity of offset responses observed (see Keller and Mrsic-Flogel, 2018). In this framework, offset responses might be due to a subtraction of sensory bottom-up signals from different predictive top-down signals in POR (i.e., “Type 2” prediction errors). Predictions of the presence of a visual stimulus, which are expected to slightly extend beyond the moment of stimulus disappearance, might explain the many types of transient offset cells (top and middle row; cf. first offset component and last three offset components in Fig. 6C). Dotted line in top row indicates that this offset response is also possible if the prediction gradually ramps across time, as the result of the rectified subtraction of stimulus signals from a predictive signal would be the same. The delayed and extended offset responses in Go and NoGo groups of cells might be generated by predictions that persist well beyond the offset of the stimulus (bottom row), similar to lateral amygdala feedback responses to POR that appear to outlast the cue duration (Burgess et al., 2016).

**Video S1. Longitudinal image registration across learning, Related to Fig. 1.**

An example field of view from two-photon calcium imaging in postrhinal cortex. The mean field of view per day from a single mouse was registered (see Methods) across 35 days of simultaneous imaging and training. The image is approximately 925 x 634 μm^2^ and has been processed by high-pass spatial filtering to deemphasize large blood vessel shadows. Blur that develops over sessions in the bottom left corner is due to bone growth.

## ACKNOWLEDGMENTS

We would like to thank Andrew Lutas, Nghia Nguyen, David Tingley, Stephen Zhang and other members of the Andermann lab for useful discussions, as well as Osama Alturkistani and Vanessa Flores-Maldonado for help performing cranial window surgeries. We would also like to thank Crista Carty and Jesseba Fernando for help with surgeries and data management. We would like to thank Alex Williams for TCA insights and GitHub support. Support was provided by an NSF Graduate Research Fellowship DGE1144152 and DGE1745303 (KLM), The Sackler Scholar Programme in Psychobiology (KLM), BBRF Young Investigator Grant (CRB), E. Matilda Ziegler Foundation for the Blind research grant (CRB), NIH F31 105678 (RNR), NIH T32 5T32DK007516 (AUS), an NIH DP2 DK105570, R01 DK109930, DP1 AT010971-02S1, R01 MH12343, a McKnight Scholar Award, a Pew Scholar Award, a Smith Family Foundation Award, a Harvard Mind Brain Behavior Interfaculty Initiative Faculty Research Award, and grants from the Klarman Family Foundation, the American Federation for Aging Research, and the Harvard Brain Science Initiative Bipolar Disorder Seed Grant, supported by Kent and Liz Dauten (MLA).

## AUTHOR CONTRIBUTIONS

KLM, CRB, and MLA designed experiments and analyses, and wrote the manuscript. KLM analyzed the data with assistance from MLA, OA, and AUS. KLM, CRB, and RNR performed two-photon imaging experiments.

## LEAD CONTACT AND MATERIALS AVAILABILITY

This study did not generate new unique reagents. Further information and requests for resources and reagents should be directed to and will be fulfilled by the Lead Contact, Mark Andermann.

## EXPERIMENTAL MODEL AND SUBJECT DETAILS

All animal care and experimental procedures were approved by the Beth Israel Deaconess Medical Center Institutional Animal Care and Use Committee. Mice (n = 5 males; n = 2 females; Emx1-Cre;Ai93 (TITL-GCaMP6f)-D;CaMK2a-tTA) were housed with standard mouse chow and water provided *ad libitum*, unless specified otherwise. Mice used for *in vivo* two-photon imaging (age at surgery: 7-17 weeks) were instrumented with a headpost and a 3 mm cranial window, centered over lateral visual association cortex including postrhinal cortex (window centered at 4.5 mm lateral and 1 mm anterior to lambda; the exact retinotopic location of visual postrhinal association cortex (POR) was determined via epifluorescence mapping; see below and Goldey et al., 2014). Portions of the data for three mice were generated as part of a previous dataset (n = 3 mice; Sugden et al., 2020). For the current study, these data were aligned longitudinally across all sessions using the methods described below (see Longitudinal image registration and ROI registration).

## METHOD DETAILS

Sample sizes were chosen to reliably measure experimental parameters, while remaining in compliance with ethical guidelines for minimizing animal use and keeping with standards in the relevant fields. Different animals in each experimental group served as replicates, and multiple repetitions of each cue also served as within-mouse replicates. Experiments did not involve experimenter-blinding, but randomization was used with respect to trial order and data collection.

### Behavioral performance and training

After at least one week of recovery post-surgery, animals were food-restricted to 85%– 90% of their free-feeding body weight. Animals were head-fixed on a 3D printed running wheel for habituation prior to any behavioral training (10 minutes to 1 hour over 3-4 days). If mice displayed any signs of stress, they were immediately removed from head-fixation, and additional habituation days were added until mice tolerated head-fixation without visible signs of stress. On the final day of habituation to head-fixation, mice were delivered Ensure (a high calorie liquid meal replacement) by hand via a syringe as part of the acclimation process. Subsequently, we trained the animals to associate licking a lickspout with delivery of Ensure, by initially triggering delivery of Ensure (5 μL, 0.0075 calories) to occur with every lick (with a minimum inter-reward-interval of 2.5 s). We tracked licking behavior via a custom 3D-printed, capacitance-based lickspout positioned directly in front of the animal’s mouth. All behavioral training was performed using MonkeyLogic (Asaad and Eskandar, 2008; Burgess et al., 2016; Livneh et al., 2017; Livneh et al., 2020; Ramesh et al., 2018; Sugden et al., 2020).

For the Go/NoGo visual discrimination task, food-restricted mice were trained to discriminate square-wave drifting gratings of different orientations (2 Hz and 0.04 cycles/degree, full-screen square-wave gratings at 80% contrast; the same 3 orientations were used for all mice; for example, during initial learning: Initially rewarded cue: 0°, Becomes rewarded cue: 270°, Unrewarded cue: 135°; grating orientations were counterbalanced across mice). All visual stimuli were designed in MATLAB and presented in pseudorandom order on a calibrated LCD monitor positioned 20 cm from the mouse’s right eye. Stimuli were presented for 2 seconds or 3 seconds (always the same for an individual mouse), followed by a 2 second window in which the mouse could respond with a lick and a 6 second inter-trial-interval (ITI). The first lick occurring during the response window triggered delivery of Ensure for the rewarded cue and delivery of quinine for the punished cue (Fig. 1B). Licking during the response window for a neutral cue did not trigger any outcome. Licking during the visual stimulus presentation was not punished, but also did not trigger delivery of Ensure or quinine. The lickspout was designed with two lick tubes (one for quinine and one for Ensure), positioned such that the tongue contacted both tubes on each lick.

Training in the Go/NoGo visual discrimination task progressed through multiple stages. The first stage was Pavlovian introduction with the Initially rewarded cue, followed by trials in which reward delivery depended on licking during the response window (“Go” trials in which licking during the response window led to delivery of 5 μL of Ensure), and finally by the introduction of punished and neutral (“NoGo” trials in which licking during the response window led to delivery of 5 μL of 0.1 mM quinine and nothing, respectively) as described in previous work (Burgess et al., 2016; Ramesh et al., 2018; Sugden et al., 2020). Imaging was performed throughout all training sessions. Mice were quickly transitioned to training involving equal numbers of each trial type, and only days in which all three cues were presented were used for subsequent data analyses. We began all imaging and behavior sessions with presentation of 2-5 Pavlovian rewarded trials (e.g., during initial learning: Initially rewarded cue; during reversal learning: Becomes rewarded cue) involving automatic delivery of Ensure reward, as a behavioral reminder. Pavlovian rewarded trials also occurred sporadically during imaging (0%– 15% of trials) to help maintain engagement. None of these Pavlovian rewarded cue presentations were included in subsequent data analyses.

Once mice stably performed the Go/NoGo visual discrimination task with high behavioral performance (d’ > 2) for at least two consecutive days, we switched the cue-outcome associations (Fig. 1F). This consisted of one of two possible “reversals” (Fig. S1A). For a type-1 reversal (n = 4/7 mice), the switching of cue-outcome associations consisted of a clockwise rotation of the outcome associated with each of the three visual oriented drifting gratings. If the initial cue-outcome associations were: Initially rewarded cue: 0°, Becomes rewarded cue: 270°, Unrewarded cue: 135°, then switching the cue-outcome associations would involve switching the Initially rewarded cue from 0° to 270°, the Becomes rewarded cue from 270° to 135°, and the Unrewarded cue from 135° to 0°. Alternatively, a type-2 reversal (n = 3/7 mice) involved swapping the neutral and rewarded cues. For example, switching the Initially rewarded cue from 0° to 270°, the Becomes rewarded cue from 270° to 0°, and the Unrewarded cue remains 135°. Immediately following reversal, more rewarded trials were given in order to facilitate learning and maintain task engagement. Over the subsequent days of the reversal learning, we quickly increased the number of unrewarded trials until equal numbers of trials of each cue type were once again presented.

Pharmacological silencing experiments have confirmed that POR, but not other areas such as secondary somatosensory cortex, are necessary for performing this visual discrimination task (Livneh et al., 2017; Ramesh et al., 2018).

### Learning stages

“Learning stages” (i.e., T1 learning, … T5 learning, T1 reversal, …, T5 reversal) were determined using the behavioral performance (*d*’) calculated for each 30-minute imaging run (1-6 imaging runs per session). Each run was binned into five evenly spaced 0.75 *d*’ bins for initial learning or reversal learning independently. Bins: *d*′ ≤ 0.75, 0.75 < *d*′ ≤ 1.5, 1.5 < *d*′ ≤ 2.25, 2.25 < *d*′ ≤ 3.0, *d*’ > 3.0. The result was 10 performance bins across the full extent of learning and relearning (Fig. 1F). While learning did progress in a largely monotonic fashion (Fig. S1B), these stages were binned by equally spaced increments in task performance rather than time.

### Mapping of retinotopic areas

To initially map the locations of lateral visual association cortex and retinotopically identified postrhinal cortex (POR), we used epifluorescence imaging of the entire 3-mm window while presenting 20° patches of noise in nine retinotopic locations. A 470-nm light-emitting diode (LED) was passed through a long-pass emission filter (500-nm cutoff), and images were recorded with an EMCCD camera (Rolera EM-C2 QImaging, 251 × 250 pixels at 4 Hz). Retinotopy was compared to Wang & Burkhalter, 2007, and the imaging field was centered over POR.

### Two-photon imaging

Two-photon calcium imaging was performed using a resonant-scanning two-photon microscope (Neurolabware; 31 frames/second; 787 × 512 pixels/frame). All imaging was performed with a 16x 0.8 NA objective (Nikon) at 1x zoom (∼1200 × 800 μm^2^). All imaged fields of view (FOV) were at a depth of 120-150 μm below the pial surface. Laser power measured below the objective was 25-60 mW using a Mai Tai DeepSee laser at 960 nm (Newport Corp.). Neurons were confirmed to be within a particular cortical area by comparison of two-photon images of surface vasculature above the imaging site with surface vasculature in widefield retinotopic maps (Andermann et al., 2011; Burgess et al., 2016; Ramesh et al., 2018; Sugden et al., 2020). For each mouse, the same field of view was tracked over multiple sessions (session count per mouse: 33, 29, 25, 23, 11, 25, 31).

## QUANTIFICATION AND STATISTICAL ANALYSIS

Statistical analyses are described in the results, figure legends, and below in this section. Statistical tests were performed using the SciPy (https://www.scipy.org/) library or using custom written software in Python for bootstrapping and permutation tests. In general, we used non-parametric statistical analyses (Wilcoxon signed-rank test for paired tests, function: scipy.stats.wilcoxon; Mann-Whitney U rank test for unpaired tests, function: scipy.stats.mannwhitneyu) or permutation tests so that we would make no assumptions about the underlying distributions of the data. All statistical analyses were performed in Python and *P* < 0.05 was considered significant (with Bonferroni correction for the number of tests, where applicable). Analyses were performed using MATLAB and Python.

### Signal extraction

We used principal/independent component analysis (PCA/ICA; Mukamel et al., 2009) and/or constrained nonnegative matrix factorization (cNMF; Pnevmatikakis et al., 2016) to extract masks of pixels with correlated activity corresponding to putative single cells. By default, we used only the top 75% of pixels with highest weights for each candidate cell mask (Ziv et al., 2013), but users screened each prospective region of interest (ROI) and could edit the size of each mask, selectively removing the lowest probability pixels. Pixels found in more than one mask were excluded from further analyses. We used a convolutional neural network to identify ROIs corresponding to putative cells, followed by user validation of each ROI. Time courses were extracted by averaging across all pixels within each binarized mask. Fluorescence time courses were extracted by unweighted averaging of the pixels within each ROI mask. Fluorescence time courses for neuropil within a 50 μm diameter annulus surrounding each ROI (but excluding adjacent ROIs and a protected ring surrounding each ROI) were also extracted (F_neuropil_(*t*): median value from the neuropil ring on each frame where *t* is the time of each imaging frame). Fluorescence time courses were calculated as F_neuropil_corrected_(*t*) = F_ROI_(*t*) – Fneuropil(*t*) + median(F_ROI_(*t*)). The change in fluorescence was calculated by subtracting a running estimate of baseline fluorescence (F_0_(*t*)) from F_neuropil_corrected_(*t*), then dividing by F_0_(*t*): ΔF/F(*t*) = (F_neuropil_corrected_(*t*) - F_0_(*t*))/ F_0_(*t*). F_0_(*t*) was estimated as the 10th percentile of a 32 second sliding window of F_neuropil_corrected_(*t*) (Livneh et al., 2020; Petreanu et al., 2012; Ramesh et al., 2018). Extracted time courses were downsampled to 15.5 Hz.

### Single-day image registration

To correct for motion along the imaged plane (x-y motion), each frame was cropped to account for edge effects (cropping removed outer ∼10% of image) and registered to an average field-of-view (cropped) using efficient subpixel registration methods (Bonin et al., 2011). Within each imaging session (one session/day), each 30-minute run (1-6 runs/session) was registered to the first run of the day. Slow drifts of the image along the z-axis were typically < 5 μm within a 30-minute run, and z-plane was adjusted between runs by eye or by comparing a running average field-of-view to an imaged volume ± 10 μm above and below our target field-of-view.

### Longitudinal image registration and ROI registration

Data registered within a day were segmented and aligned in two rounds to maximize the number of cells that were captured consistently across days while minimizing laborious steps of removing “false positive” cells. Probabilistic cross-day region of interest (ROI) alignment was accomplished using the CellReg algorithm (Sheintuch et al., 2017) in combination with custom MATLAB image registration software relying on displacement field estimation using the Demon’s algorithm (Thirion, 1998; Vercauteren et al., 2009; MATLAB 2017b functions: imregdemons and imwarp). We first tested this cross-day alignment on the outputs of field-standard cell segmentation algorithms relying on PCA/ICA (Mukamel et al., 2009) or constrained nonnegative matrix factorization (cNMF; Pnevmatikakis et al., 2016). We found that PCA/ICA mask positions were more consistently registerable across days, while many cNMF mask positions were less consistently registerable compared to our own “ground truth” alignment of the same data (see Ramesh et al., 2018, for details). However, cNMF produced a much larger number of masks. While cNMF often produced many ROIs in a given session that were not possible to align to any other imaging session, it also often identified masks that were well aligned to masks from other sessions, despite the fact that masks could exhibit little to no meaningful temporal activity. This was important for the current study, as we aimed to track cells across many days, even on days in which they were not active. By combining both algorithms in series, relying primarily on PCA/ICA, but then using cNMF to systematically fill in the “holes” in alignment across days in already aligned and user-validated PCA/ICA segmentation masks, we took advantage of the benefits of both algorithms. Steps for registration, segmentation, and cross-day alignment process are described in more detail below.

First, mean images from each day of imaging were registered within day using custom MATLAB software performing rigid body registration (see Single-day image registration; as in Ramesh et al., 2018). The mean registered field of view per day was then registered to each additional daily field of view using custom MATLAB image registration software relying on displacement field estimation using the Demon’s algorithm (Thirion, 1998; Vercauteren et al., 2009; MATLAB 2017b functions: imregdemons and imwarp) where the field of view from one day was defined as the registration target. From this, the best registration set was defined (i.e., the registration target day with the most successfully matched fields of view across other days) and any days that were not possible to register were excluded from further analysis. An example of the resulting alignment across days is shown in Video S1. While registration to the standard reference target described above failed on some days, these days could sometimes be successfully aligned to another day within the set (e.g., if day 2, 3, and 4 could be registered to day 1, but day 5 could only be registered to day 4). In this case, sequential registration was used (e.g., day 5 was first aligned to day 4 and then the registration mapping day 4 to day 1 was also applied). This allowed us to maximize the number of daily sessions aligned across training and imaging. These sets of warp fields could later be applied to register segmentation masks across days of imaging.

The binarized masks identified using an initial round of PCA/ICA segmentation (see Signal extraction) across days of imaging were then probabilistically registered using the CellReg algorithm (Sheintuch et al., 2017). This produced centroids of each cell’s position across registered days. These centroids were then used to “seed” (i.e., initialize) nonnegative matrix factorization which resulted in another set of prospective regions of interest which were screened initially using a convolutional neural network and later by a user. For this set of validated cNMF masks we identified putative novel masks by comparing binarized masks for each session, subtracting PCA/ICA masks from cNMF masks. cNMF masks that were less than 8 pixels or were “hole punched” by this operation were removed. The remaining cNMF masks were considered putative ROIs that the original round of PCA/ICA segmentation had missed.

We once again repeated probabilistic registration using the CellReg algorithm except now we pooled masks identified using PCA/ICA or cNMF. cNMF masks that were registered to only other cNMF masks were removed from analysis, while the remaining masks represented ROIs that had now filled in a “hole” in our original PCA/ICA segmentation. From these binarized masks signal extraction was performed. For details on signal extraction, see Signal Extraction.

For 3/7 mice (existing datasets from Sugden et al., 2020) with segmentation and signal extraction already complete, cells were aligned using the same process as outlined above, but no additional step was taken to identify additional ROIs using cNMF. Instead, mean fields of view were registered across days, this registration was applied to the masks for each identified ROI, and the CellReg algorithm (Sheintuch et al., 2017) was used to align and probabilistically register cells across days of imaging. While this likely results in cells tracked over a smaller proportion of total sessions in these 3 mice, we did not observe any systematic differences in results from these 3 mice versus the other four mice for which PCA/ICA followed by cNMF was used, and thus all datasets were pooled.

In all cases, cells with a registration score of less than 0.8 were excluded from further validation and analysis. Of the 8303 total cells registered across 7 mice, 7824 met this criterion and were included in subsequent analysis.

### Offset responsiveness

To detect and separate putative offset and onset cells prior to determining whether such cells are significantly driven (see Cell drivenness section, below), we calculated the slope of the average stimulus-response (averaged across fluorescence traces from all trials; z-scored Δ*F*/*F*) during the one second prior to stimulus offset. This slope was subtracted from the average stimulus-response waveform. We then calculated a new slope from this slope-subtracted response trace, between (i) the offset of the stimulus and (ii) the first subsequent time point for which the offset response crossed 75% of its maximum response following the offset of the stimulus. Cells were defined as preferentially offset responsive cells if two conditions were met: 1) the peak of the waveform occurred at least 120 milliseconds after stimulus offset, to account for slow dynamics of GCaMP6f related to activity occurring prior to cue offset; 2) the angle of the slope (after subtraction) estimated for the period following the stimulus (i.e., time of offset to time of first crossing of 75% of peak response) offset was larger than 0.06°. Cells that did not pass these two conditions were defined as potential onset responsive cells. Both preferentially offset responsive cells and onset responsive cells were later assessed as to whether they were significantly driven for onset or offset responses respectively.

### Cell drivenness

We determined whether a cell was driven (i.e., whether it demonstrated statistically significant responses to the onset or offset a given stimulus type during a given stage of learning) as follows. For each stage of learning, we assessed whether a neuron had a significant increase in GCaMP6f fluorescence (z-scored Δ*F*/*F*) in at least one of two windows during the cue presentation, relative to baseline (defined as −1 to 0 s before cue onset). For responses to the stimulus onset, the baseline was compared to the average activity across the entire stimulus period (0 – 2 s or 0 – 3 s duration), and was also compared to the average activity across a 500-millisecond window from 200 to 700 milliseconds following stimulus onset (to ensure that short-latency, transient responses common in initial inspection were not unfairly excluded). Significantly driven responses were estimated using a one-way Wilcoxon signed-rank, Bonferroni-corrected by 2 to account for the number of windows of time over which comparisons were calculated. To determine stimulus offset responsivity, a similar process was repeated, but two baseline periods were used: (i) −1 – 0 seconds before stimulus onset and 2) −1 – 0 seconds before stimulus *offset*. The average activity for each baseline was compared to the average activity across the entire response window (0 – 2 seconds after stimulus offset) as well as a 500-millisecond window from 200 to 700 milliseconds following stimulus offset. Each of these two baseline comparisons were Bonferroni-corrected for both windows of comparison independently. For a cell to be considered significantly offset responsive, it had to have a response that was significantly above the pre-stimulus baseline as well as rise significantly at the offset of the visual stimulus. We used a conservative criterion for significantly responsive cells: *P* < 1.3 × 10^-4^. We later confirmed that our results held with different thresholds of responsiveness, but we found this stringent criterion eliminated many “noisy” and inconsistent cells with temporal dynamics that would otherwise be difficult to estimate precisely.

### Average stage response traces

Many analyses and visualizations were performed using traces that were the average responses per stage for each cell including a 1 second baseline window and a 2 second stimulus window (or reward response window, for preferentially offset responsive neurons) concatenated across stages per cell. The process of calculating these traces is described in more detail below.

Cells were initially divided into groups that responded preferentially during cue onset or following cue offset (see Offset responsiveness) and these two groups were treated independently for all data organization steps.

To estimate the average response per stage for each cue presentation, extracted traces (see Signal extraction) were first concatenated across 30-minute imaging runs and z-scored. From this z-scored trace, each trial, including a 1 second baseline window prior to stimulus onset (or prior to stimulus offset for offset responsive neurons) and a 2 second window starting at stimulus onset (or following stimulus offset for offset responsive neurons), was selected and re-zeroed by subtracting the mean activity in the 1 second prior to visual stimulus onset. This was also true of offset cells where the mean of the 1 second baseline period *prior* to visual stimulus *onset* was subtracted from each trace. The average response per stage for each cue type was then calculated by first pooling trials for each day and averaging and subsequently pooling days for each stage of training and averaging. For each cell, the average activity for each stage was then concatenated into a single vector with the mean response in each of the ten stages of training arranged sequentially. Responses for each cell were then normalized to the maximum response across all stages and cues (i.e., “peak normalized”). Data were organized in the form [neurons] x [peak-normalized concatenation of stage-averaged time courses for each neuron] x [“Initially rewarded” cue, “Becomes rewarded” cue, “Unrewarded” cue].

Cells were assessed for their drivenness during each stage of learning independently (see Cell drivenness) and cells were included if (i) they were significantly driven on at least one stage of training for at least one of the three cues, (ii) if they had segmentation masks on two or more stages of training, and (iii) if they had non-suppressed responses (i.e., all averaged evoked responses were not exclusively negative) to either to stimulus onset (or to stimulus offset for preferentially offset responsive neurons). Of the 7824 cells that were registered with high confidence across 7 mice, 2244 passed these additional criteria. This operation was repeated for each cell across all mice.

### Tensor Component Analysis

Tensor component analysis (TCA) was fit using the *tensortools* package (Williams et al., 2018) in Python. In all analyses, nonnegative CP Decomposition using the Hierarchical Alternating Least Squares (HALS) method was used. We found this method to be especially valuable because missing values (e.g., cells for which a cell mask was not available for a given stage of learning) can be masked and updated during fitting, while removing their contribution from the calculation of the cost function. This allowed us to use a much larger dataset, enabling cell classification based on across-stage response dynamics that would likely be less tractable using other dimensionality reduction and embedding techniques.

For TCA, all mean response time courses for each of the three cues, for all stages for which data was available, were pooled across mice to form a three-dimensional matrix or “tensor” (dimensions: cells *x* stages *x* cue types). This final tensor was used for TCA fitting.

Tensor component analysis requires empirical fitting to determine an appropriate rank (i.e., number of components) to represent the data. Accordingly, we fit TCA from rank 1-20, 40, and 80 and evaluated the model reconstruction error over these ranks to determine where an approximate “knee” or “elbow” existed. We determined that ranks 6-12 for onset responsive cells and ranks 5-11 of offset responsive cells fell within a reasonable range of model performance without excessive reduction in performance relative to higher model ranks. For each rank, we randomly initialized and fit independent models 10 times and compared the quantitative (see Williams et al., 2018) and qualitative similarity across models. The ranks selected were confirmed to have similarity scores above what is considered appropriate in Williams et al., 2018 (i.e., similarity ≥ 0.8). We chose to select models central to the ranges we had defined for further analysis (i.e., rank 9 for onset responsive neurons and rank 8 for preferentially offset responsive neurons), but it is important to note that even across other nearby ranks, models produced similar results (see Figs. S2 and S6 for additional details).

### Clustering cells using TCA

Tensor component analysis was used to cluster cells based on the cell factor weights of each component after “absorbing” the weights of the temporal factor weights and tuning factor weights into the cell factor weights by rescaling. Specifically, since each TCA component is the outer product of each of its constituent factors (here, [cell factor weights] * [temporal factor weights] * [cue tuning factor weights]), each component can be described equivalently using, for example, cell factor weights that are 2x larger, together with temporal factor weights that are 2x smaller (i.e., a single component = [cell factor] * [temporal factor] * [tuning factor] = 1 * 1 * 1 = 2 * 1 * 0.5). Thus, we were able to define interpretable clusters by rescaling temporal factors and tuning factors to 1 (dividing by the maximum of each factor) and adjusting the cell factor weights accordingly (i.e., by multiplication by the scale factors that were used to normalize the temporal and tuning factors). This meant that temporal factors were now the same scale as the input data, since the response traces concatenated across stages had a maximum response value of 1. After rescaling, tuning factors also had a maximum value of 1 and empirically, tuning factor vectors usually had values near 0 and 1, across all three cues. In this way, cell factors could then be interpreted as groups of cells that shared a particular type of activity (with common within- and across-stage response dynamics and cue tuning) across stages of training. Component “clusters” of cells were determined by assigning each cell to the component for which they had the highest rescaled cell factor weight.

### Analysis of possible epileptiform events

The Emx1-Cre;Ai93;CaMK2a-ttA mouse line (Madisen et al., 2015) used in our work has been reported to exhibit epileptiform events in some cases (Steinmetz et al., 2017). We confirmed that back-crossed mice from this strain did not exhibit pathological activity (see below and Sugden et al., 2020). Early experiments in our lab using mice obtained directly from the Allen Brain Institute did appear to exhibit visible epileptiform events as detected in the neuropil signal in cortical imaging. Those mice were excluded from this study. We then acquired mice from Jackson labs (line 024108 rather than 024103 used in Steinmetz et al., 2017) which may have been further back-crossed. To further address the possibilities that epileptiform events occurred in the 7 mice used in our study, we characterized the amplitude and width of all transient events in the neuropil from spontaneous activity recordings in all of our mice (as in Steinmetz et al., 2017). We plotted peak width versus event peak prominence (height above local background) as in Steinmetz et al., 2017 and Sugden et al., 2020. We found that 6/7 mice did not show any evidence of epileptiform events, while 1/7 mice only demonstrated a small number of possible epileptiform events on a single day of imaging. This day of imaging was excluded from further analysis.

### Onset/offset latency and slope

Onset (offset) latency was calculated as the time to first crossing of a threshold (30% of peak response) following stimulus onset (offset) (Figs. 3B, 7B) for averaged traces across all cells in a component cluster (one cluster was defined for each TCA component and mouse, see above). This was calculated as follows. For each component cluster and each mouse, the average activity per stage was calculated across cells from their average response trace, using the preferred tuning of each cluster and excluding undriven stages per cell, resulting in a component cluster average across stages. Responses were then pooled across stages and averaged so that the average activity across learning for each component cluster was represented by a single trace comprised of the 1 second baseline and the 2 second window following stimulus onset. For offset responsive cells, the last second of the stimulus and the 2 second post-stimulus window were used. This 3-second trace was then upsampled from 15.5 Hz to 1000 Hz using linear interpolation (function: scipy.interpolate.interp1d). For onset cells, the response latency was calculated as the time to first crossing of a value of 30% of the peak response following stimulus onset. For offset cells, the same calculation was made after first re-zeroing the trace by subtracting the mean of last second of the stimulus. Peak response latency (used for sorting in Figs. 1G and 6B) was calculated as the time to the peak (i.e., 100%) response.

For estimation of onset or offset slopes, the process was the same as above, but instead of the latency, we calculated the slope between 30% and 80% of the peak response following stimulus onset or offset, respectively (Figs. 3B, 7B).

### Sustainedness index

We estimated whether the time course of the response to a cue onset (or offset) was either adapting or showed sustained activity late in the stimulus presentation by defining a sustainedness index for each component cluster and each mouse was calculated as follows. The average activity per stage was calculated across cells in a component cluster, using the preferred tuning of the TCA component associated with the cluster, and excluding undriven stages per cell, resulting in a component cluster average for each stage. Stages were then averaged so that the average activity across learning for each component cluster was represented by a single trace with from 1 second prior to 2 seconds after stimulus onset (for onset responsive cells) or from 1 second prior to 2 seconds after stimulus offset (for offset responsive cells). For onset cells, sustainedness index was calculated using an estimate of the “early” response to stimulus presentation (“Mean_early_,” from 65 to 565 milliseconds after stimulus onset), and an estimate of the “late” response to stimulus presentation (“Mean_late_,” 1500-2000 milliseconds after stimulus onset). Similarly, “early” and “late” offset responses were calculated using the 65-565 milliseconds and 1500-2000 milliseconds windows following stimulus offset. The sustainedness index was calculated as: Sustainedness index = [Mean_late_ - Mean_early_] / [Mean_late_ + Mean_early_] (Figs. S3A, S7A). Thus, a component cluster with a highly sustained response (e.g., elevated activity during late periods after stimulus onset or offset, but not during periods soon after onset or offset), would have a value close to 1; clusters with a short-latency, transient response would have a value close to −1; and clusters with a perfectly flat response beginning soon after cue onset would have a sustainedness index of 0.

### Peak stage response

We estimated the stage with the strongest response for the mean traces for each component cluster and mouse, as follows. The mean evoked response for each stage was computed over the 2 second period following stimulus onset or stimulus offset (after baseline subtraction), yielding a vector of response values, one for each of the ten stages (i.e., one vector for each component cluster, with a single value for each stage from T1 learning through T5 reversal). The peak stage was calculated as the stage with the maximal response (Figs. 3B, 7B).

### Cue preference

Cue preference for each cell was determined using the cue tuning factors for the TCA component for which a cell had the largest cell factor (i.e., the component cluster to which the cell belonged). Cue tuning factor weights, which scaled between 0 and 1 (Fig. 2C), were used to determine cue preference. A cell was determined to “prefer” a given cue if its tuning factor weight for that cue was greater than 0.5, and less than 0.5 for the other two cues. While this threshold value was arbitrary, all thresholds tested yielded similar results since tuning factors took values that were almost exclusively 1 or 0 for each cue (Fig. 2C). Cells were considered to prefer the “Initially rewarded” cue, “Becomes rewarded” cue, or “Unrewarded” cue if the tuning factor had only a single value greater than 0.5 to one of these three cues. Cells were considered “broadly” tuned if tuning factor values were above 0.5 for all cues. Cells were considered “joint” tuned if two out of the three cue values were above 0.5 (e.g., Fig. 6C; 4^th^ and 5^th^ rows). For all subsequent calculations, unless otherwise stated, all trials for each of a cell’s “preferred” cue types (i.e., for all cues in which the cue tuning factor weight was greater than 0.5) were used.

### Go modulation

Go modulation was defined for each cell and learning stage as the modulation of responses to a cue depending on whether the animal licked (“Go trial”) during the 2 second post-stimulus window (the operant response window) or didn’t lick during that window (“NoGo trials”). First, we calculated a single response value per trial, averaged over for the first 2 seconds of the stimulus response (for onset response analyses) or the first 2 seconds following offset (for offset response analyses). Average Go and NoGo responses were then calculated as the mean responses across Go trials or NoGo trials, respectively, averaged within each day then averaged across days within each stage. The two vectors of average Go or NoGo responses per stage were then normalized using a single value — the peak response across stages estimated using the maximum response across Go and NoGo responses. Peak-normalized Go and NoGo responses were then separately averaged across stages (excluding stages where that cell was not driven; see Cell drivenness) to obtain a single value for each cell. The Go modulation index was calculated as: average driven Go response minus the average driven NoGo response.

### Running modulation

Running and stationary trials were determined using the average running speed in the 1 second preceding stimulus onset. Running trials were defined as those trials where an animal’s speed was > 10 cm/s, and stationary trials were defined as those trials where an animal’s speed was < 4 cm/s. To calculate running modulation, we first calculated a single response value per trial, averaged over for the first 2 seconds of the stimulus response (for onset response analyses) or the first 2 seconds following offset (for offset response analyses). Average running and stationary responses were then calculated as the mean responses across running trials or stationary trials, respectively, first averaged within each day then averaged across days within each stage. This vector of average values per stage for running and stationary trials was then normalized to the peak response across stages using the maximum response pooling running and stationary responses. The Running modulation index was calculated per stage as: average running response minus the average stationary response. For calculations that pool across stages, peak normalized running and stationary responses were then separately averaged across stages (excluding stages where that cell was not driven, see Cell drivenness) to obtain a single value for each cell before calculating the Running modulation index as above. To estimate the stability of running modulation across stages, we calculated the running modulation stability index (RMSI) as RMSI = [(number of stages in which cue response during running trials was greater than during stationary trials) - (number of stages in which cue response during stationary trials was greater than during running trials)] / (total number of stages where the cell was driven by the cue). Cells with an RMSI of −1 consistently preferred stationary trials, while cells with an RMSI of 1 consistently preferred running trials across driven stages of training.

### Support vector machine trial classification

To determine whether we could classify cue identity on stationary trials when training on data from running trials and vice versa (Fig. 4H), we used a support vector machine (SVM). The SVM was trained using sklearn.svm.SVC from the scikit-learn (scikit-learn.org) API in Python. For each day, an SVM was trained on 80% of running trials, and tested on a held-out set of 20% of running trials as well as a held-out set of all stationary trials. This process was repeated using stationary trials as the training set. A balanced number of trials of each cue type were used for training (i.e., stratification). For each locomotor context (running trials or stationary trials), models were trained and the average performance (e.g., fraction of correctly classified trials) across days was calculated for each mouse (Fig. 4G).

### Population Variation

Population variation was used to calculate the variability in the average population activity (z-scored Δ*F*/*F*) across stages of learning (Fig. 5G,K). First, we calculated the average response per stage per cell, by taking the average evoked response across the 2-second stimulus window, resulting in a vector for each cell with a single value per stage. We then calculated the population response by averaging across all cells for each stage. Finally, we quantified the population variation by computing the standard deviation in the population response across stages. We sought to test if the relative stability in the average magnitude across all neurons that we observed across stages of learning (Fig. 5A,C,E) was simply due to chance (e.g., cause by averaging many cells together) or if this was unexpected and due to a specific balancing of the responses of different component clusters of cells at each stage of learning. Accordingly, we shuffled the identity of learning stages separately for each cluster of cells, and then recalculated the population variation. We repeated this procedure 10,000 times to obtain a distribution of standard deviation values that could have been expected by chance over clusters of neurons.

### Bootstrapping confidence intervals across components

For each mouse, values (e.g., Go modulation) were shuffled across components with replacement, and the mean value was calculated for a “random” component cluster. This process was repeated 100,000 times to obtain a distribution of mean values for a component cluster across mice that could be expected by chance. Confidence intervals, alpha=0.05 (i.e., percentiles 0.025 and 0.975), were drawn onto the plot. If the plot includes a datapoint for “All cells,” this point was excluded from the shuffle, as the shuffle was meant to test how likely it is to get a component cluster with an unexpectedly positive or negative value given random assignment of component cluster identity during the calculation.

### Bootstrapping statistic across stages of learning

For comparisons between stages of learning (e.g., T4 versus T5 learning), we performed the same calculation comparing the average peak-normalized values between two stages of learning after shuffling the identity of stages to test the possibility that the difference calculated between the stages of learning of interest could have occurred by chance. We re-ran this shuffle 10,000 times and compared the real data to the distribution of shuffled values. Tests were Bonferroni corrected for multiple comparisons, which included a test for each component and the “All cells” test (e.g., Figs. 3G, S7C).

### Bootstrapping statistic across contexts

For comparisons between contexts, the exact same calculation was performed (i.e., Running modulation or Go modulation), but for each cell, the identity of running trials and stationary trials (or Go and NoGo trials) was randomized per stage before calculating Running modulation (or Go modulation) for each component cluster. This process was repeated 10,000 times to assess the possibility that the Running modulation (or Go modulation) would have been produced by chance from the underlying data. Tests were Bonferroni corrected for multiple comparisons, which included a test for each component and the “All cells” test.

## DATA AND CODE AVAILABILITY

The datasets and code are available upon request to the Lead Contact.

## KEY RESOURCE TABLE

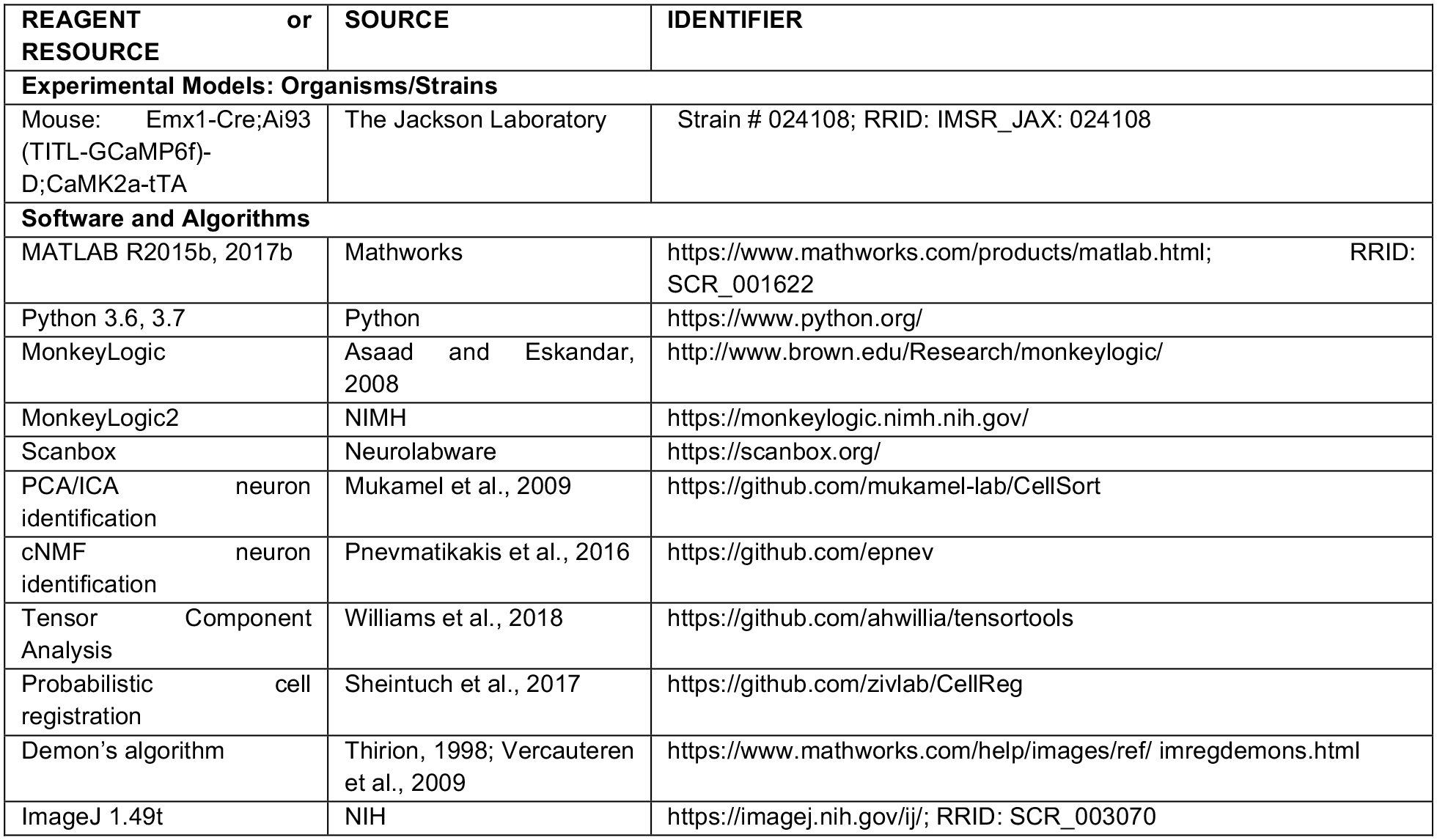

